# The non-coding RNA miR-17~92 is a central mediator of T cell activation

**DOI:** 10.1101/2020.10.13.336537

**Authors:** Marianne Dölz, John D. Gagnon, Mara Kornete, Romina Marone, Glenn Bantug, Robin Kageyama, Christoph Hess, K. Mark Ansel, Denis Seyres, Julien Roux, Lukas T. Jeker

## Abstract

T cell activation is paramount for productive adaptive immune responses. CD28 is a key clinically targeted immunoregulatory receptor because it provides the prototypical costimulatory signal required for T cell activation. Therefore, a precise understanding of the molecular consequences of CD28 costimulation has direct therapeutic relevance. Here, we uncover that the microRNA cluster miR-17~92 is part of the molecular program triggered by CD28 costimulation and hence T cell activation. Combining genetics, transcriptomics, bioinformatics and biochemical miRNA:mRNA interaction maps we demonstrate that transgenic miR-17~92 can cell-intrinsically largely overcome defects caused by CD28-deficiency. miR-17~92 promotes transcription of a proinflammatory gene signature by enhancing the calcineurin/NFAT pathway. miR-17~92 binds to and represses a network of genes including several negative regulators of T cell activation. Finally, CD28-deficient T cells exhibit derepressed miR-17~92 target genes during activation, demonstrating that this non-coding RNA is required to shape the transcriptome. Thus, we propose that miR-17~92 constitutes a central mediator for T cell activation, integrating signals by the TCR and CD28 costimulation. In this model miR-17~92 facilitates T cell activation by dampening the breaks that prevent T cell activation.

## Introduction

T cells are paramount to protect mammals from infections and tumors. T cell activation, a key event for adaptive immunity, relies on two signals: T cell receptor (TCR) stimulation as well as costimulation by specialized receptors. While the TCR signal provides specificity, costimulation by antigen presenting cells (APCs) provides quantitiative and qualitative support for T cell activation ^1,2^. One of the best studied and prototypical costimulatory molecules is CD28. It promotes multiple processes required for T cell biology such as T cell activation, proliferation, survival, metabolic adaptation and Interleukin-2 (IL-2) production ^2,3^. These form the basis for clonal expansion and T cell differentiation into a variety of effector T cells that are necessary to mount effective immune responses. Recent data further demonstrates that CD28 expression is not only required for T cell priming but also days later for effector CD4^+^ T cell responses during infection ^4^.

Due to its centrality for T cell and immune responses more generally, CD28 is an important target in therapeutic immunology ^2,5,6^. CD28 blockade by CTLA4-Ig is clinically used to prevent renal allograft rejection and to treat rheumatic disease ^2^. In contrast, vaccine adjuvants induce activation of innate immune cells to express ligands that trigger costimulatory molecules on T cells. Furthermore, immune-activating CTLA-4 blocking antibodies represent the foundation of cancer immunotherapy ^7^ and CD28 is required for cancer immunotherapy with PD-1 blocking antibodies ^8–10^. Finally, CD28 intracellular signaling domains can provide the required costimulatory signal in 2^nd^ and 3^rd^ generation chimeric antigen receptor (CAR) T cell constructs for adoptive cellular therapies ^11^. Thus, CD28 is a key clinically relevant immunoregulatory receptor and a precise understanding of molecular events induced by CD28 costimulation has direct therapeutic relevance. However, despite intense research, the understanding of the molecular consequences of CD28 costimulation remains incomplete ^2^. CD28 costimulation acts through pleiotropic effects; it promotes the phosphatidylinositol 3-kinase (PI3K) pathway, amplifies the TCR signal, stabilizes the transcriptome induced by TCR stimulation and enhances calcineurin/NFAT signaling ^2,3,12^. Collectively, CD28 stimulation ultimately leads to transcriptional changes mediated by the transcription factors (TF) NF-κB, AP-1 and NFAT ^2^. In addition, CD28 also acts through many non-transcriptional mechanisms such as mRNA stabilization and altered mRNA splicing ^2^. Thus, due to the complexity of CD28 costimulation, substantial controversy remains about the importance of various molecular mechanisms and the relative importance of quantitative versus qualitative CD28-mediated signals ^1–3,5^.

Since a protein-centric view prevailed in most studies, here we investigated the role of the microRNA cluster miR-17~92, a non-coding RNA, in CD28 costimulation and T cell activation. MicroRNAs (miRNA) are arguably the best studied class of non-coding genes. These short RNA molecules of ~22nt length are highly conserved gene repressors that mainly act through base pairing with the 3’ untranslated region (UTR) of target RNAs resulting in their decreased abundance and/or translational inhibition ^13^. Together with TFs, miRNAs are the most important trans-regulators of gene expression, regulating most mRNAs ^14^. Target RNAs can be bound by multiple miRNAs and individual miRNAs can bind to and repress multiple genes, often genes found in the same pathway ^13^. Canonical base-pairing is largely determined by miRNA nucleotide positions 2-7, called the “seed” region ^14^ but non-canonical miRNA targeting is widespread and can be equally effective ^15,16^. The interaction of a miRNA and its target mostly results in mild gene repression, often only reducing protein concentration by less than 2-fold. Nevertheless, miRNA-mediated gene regulation is highly consequential for T cell differentiation and function ^13,17–19^.

We noticed that key functions attributed to CD28 such as T cell proliferation, survival and T_FH_ differentiation are also regulated by miR-17~92. miR-17~92, a polycistronic transcript induced by CD28 costimulation ^20^, gets processed into 6 mature miRNAs representing 4 different seed families ^17^. Much like CD28, miR-17~92 promotes T cell proliferation, survival and differentiation ^19,21^. Moreover, T cell-specific miR-17~92-deficiency results in severely impaired T_FH_ differentiation and impaired GC formation ^22,23^, reminiscent of impaired GC formation observed in CD28-deficient mice ^24^. On the contrary, miR-17~92 overexpression results in a systemic lupus-like syndrome with increased GC formation and autoantibody production ^21^ most likely due to enhanced T_FH_ generation ^22,23^.

Here, we show that miR-17~92 was necessary to mediate part of the molecular changes induced by CD28 costimulation and its forced expression was sufficient to restore an important fraction of the impaired transcriptional regulation and function of CD28-deficiency in murine CD4^+^ T cells. We used transcriptome analysis, computational predictions and biochemical miRNA/target RNA interaction maps to define a high confidence set of 68 empirically validated direct miR-17~92 target genes in T cells. Pathway analysis revealed that miR-17~92 promoted expression of NFAT-regulated genes. Pharmacologic experiments using the calcineurin inhibitor cyclosporin A (CyA) confirmed that miR-17~92 promoted the calcineurin/NFAT pathway. Our results identify miR-17~92 as an important mediator of CD28-costimulation induced T cell activation that promotes key activating pathways. We propose that miR-17~92 represses multiple inhibitory proteins that restrain T cells from becoming activated.

## Results

### miR-í7~92 regulates T cell proliferation and Interleukin-2 production

In order to further investigate the connection between CD28 and miR-17~92, we analyzed the consequences of T cell-specific loss and gain of miR-17~92 on key processes regulated by CD28. We compared samples from mice that lack miR-17~92 in T cells (CD4cre.miR-17~92^lox/lox^, designated T^1792Δ/Δ^ hereafter), wildtype (wt) mice and mice overexpressing miR-17~92 in T cells (CD4cre.Rosa26^lox^STOP^lox^CAG-miR-17~92Tg, designated T^1792tg/tg^ hereafter). Confirming previous findings, we showed that miR-17~92 promotes T cell proliferation (Fig. 1a) ^22,23,25^. In addition, in comparison to wt T cells, production of the classic CD28-dependent gene interleukin-2 (IL-2) was impaired in T^1792Δ/Δ^ T cells and increased in T^1792tg/tg^ T cells. Intracellular IL-2 and IL-2 secretion correlated with the miR-17~92 genotype (Fig. 1b, c). Thus, miR-17~92-deficiency was reminiscent of phenotypes observed in CD28-deficient mice. In contrast, transgenic miR-17~92 appeared to enhance T cell activation/costimulation. Thus, the data suggested that the non-coding RNA miR-17~92 exerted a dose-dependent regulation of CD4^+^ T cell proliferation, IL-2 production, and more complex biologic functions such as T_FH_ differentiation^22,23^, all of which are known to be CD28-dependent.

**Figure 1.**
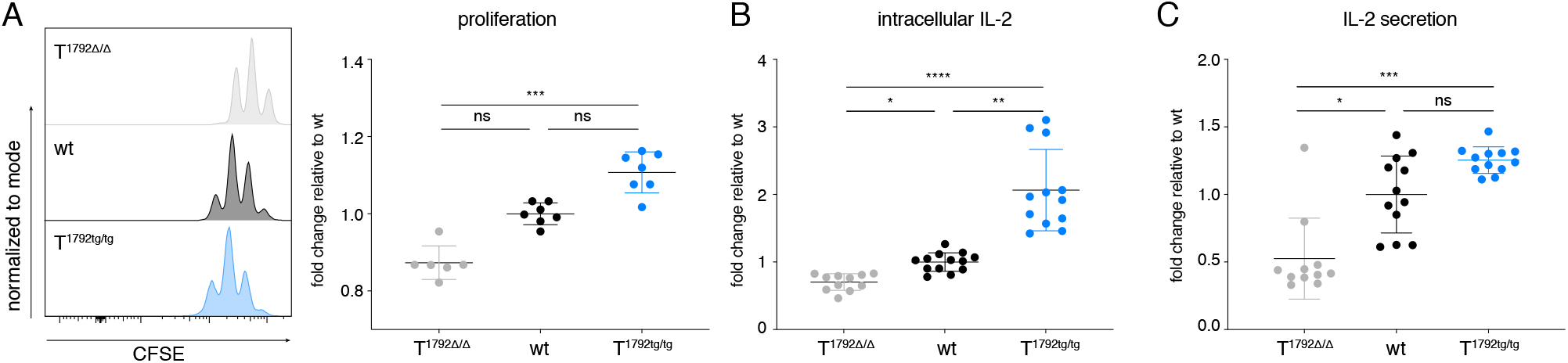
miR-17~92 regulates T cell proliferation and Interleukin-2 production. A) Proliferation of CFSE labelled T1792Δ/Δ (grey), wt (black) and T1792tg/tg (light blue) CD4+ T cells activated for 48h with plate-bound αCD3/αCD28 mAbs. B) Quantification of flow cytometric intracellular IL-2 staining in CD4+ cells stimulated for 3h with PMA/Iono/BFA. C) IL-2 secretion measured by ELISA in culture supernatants at 48h. Error bars show mean ±SD, Dunn’s multiple comparison test, p values: ns=not significant, *<0.05 **<0.002 ***<0.0002 ****<0.0001. Data represent 2-3 independent experiments with 3-4 biological replicates per group.

### Transgenic miR-17~92 enables T cell activation of CD28-deficient T cells in vitro

Since miR-17~92 expression is induced in a CD28-dependent manner during T cell activation ^20^, we hypothesized that miR-17~92 could be a downstream mediator of CD28 costimulation. To test this hypothesis we crossed B6.CD28^-/-^ (CD28^-/-^) mice ^26^ with T^1792tg/tg^ mice, resulting in B6.CD28^-/-^.CD4cre.Rosa26^lox^STOP^lox^CAG-miR-17~92Tg, designated “rescue” hereafter. T cells in these mice lack CD28 but constitutively express transgenic miR-17~92. If miR-17~92 physiologically transmitted signals induced by CD28 then we expected that transgenic miR-17~92 expression could restore some of the CD28 defects. First, we investigated how “rescue” cells behaved in vitro. We compared wt, CD28^-/-^ and “rescue” naïve CD4^+^ T cells stimulated with plate-bound anti-CD3 mAb alone to mimic signal 1 or a combination of anti-CD3 and anti-CD28 mAb to mimic combined signals 1 and 2. As expected, wt cells blasted (Fig. 2a) and increased proliferation (Fig. 2b) in response to costimulation compared to anti-CD3 stimulation alone. Compared to wt cells, CD28^-/-^ T cells showed reduced size and proliferation and were unable to respond to anti-CD28 stimulation (Fig. 2a,b). In contrast, “rescue” T cells stimulated with anti-CD3 alone or combined anti-CD3/anti-CD28 blasted and proliferated like wt T cells fully stimulated with anti-CD3 and anti-CD28 mAb (Fig. 2a, b). Since IL-2 production is a major consequence of CD28 costimulation and miR-17~92 correlated strongly with IL-2 production (Fig. 1b,c) we analyzed this cytokine next. As with blasting or proliferation, CD28-deficient T cells produced less IL-2 but “rescue” T cells produced even supraphysiologic amounts of IL-2 (Fig. 2c). Thus, transgenic miR-17~92 was sufficient to replace CD28 for three major costimulation-dependent processes and IL-2 was very sensitive to miR-17~92 expression. Since these experiments were carried out in the presence of exogenous IL-2 it is unlikely that increased blasting and proliferation were a consequence of increased IL-2 but rather reflected a cell-intrinsic effect.

**Figure 2.**
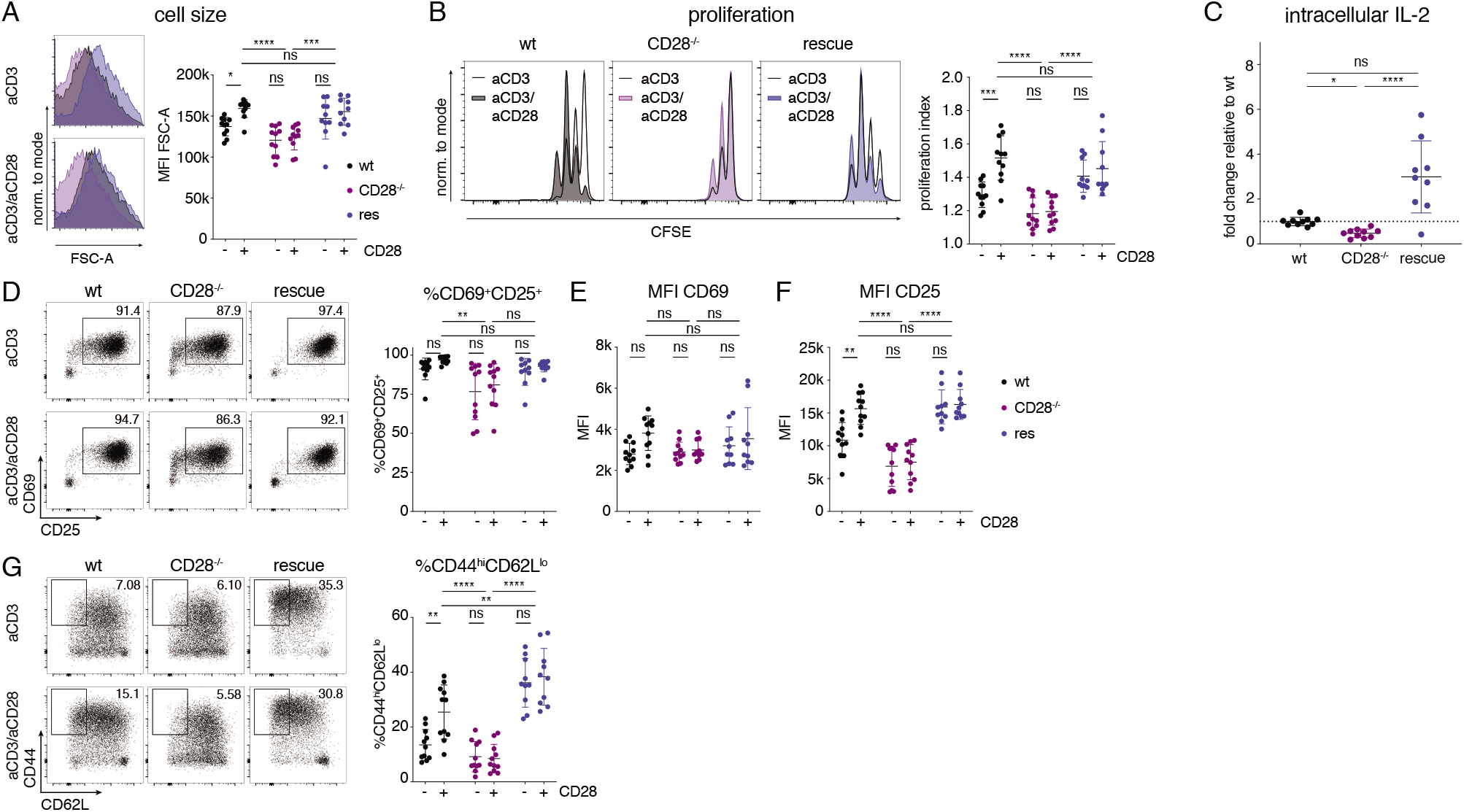
Transgenic miR-17~92 enables T cell activation of CD28-deficient T cells in vitro. wt (black), CD28-/- (purple), rescue (dark blue). A) Blasting of CD4+ T cells shown as MFI of FSC-A of the lymphocyte gate. B) Proliferation measured by CFSE dilution, gated on viable CD4+ T cells. Representative histograms of each genotype activated without (blank) or with (colored) αCD28. C) Quantification of flow cytometric intracellular IL-2 staining in CD4+ cells stimulated for 3h with PMA/Iono/BFA. D-G) CD4+ T cells were stimulated for 48h with plate-bound αCD3 with (+) or without (-) αCD28 and investigated for expression of early activation markers CD25/CD69 expression (D, E, F) as well as CD44/CD62L expression (G). Data from 3 independent experiments with 3-4 biological replicates per group. Error bars represent mean ±SD, Tukey’s or Hom-Sidak’s multiple comparison test; p values: ns=not significant, *<0.05 **<0.002 ***<0.0002 ****<0.0001

Next we turned our attention to surface markers whose expression is either TCR- or CD28-dependent. The early activation marker CD69 is rapidly upregulated during T cell activation and reflects TCR signaling strength ^27^. As a second marker we used the high affinity IL-2 receptor alpha subunit CD25, a well-known CD28-dependent gene. The relative number of cells that expressed CD69 and CD25 (Fig. 2d) and CD69 expression per cell (Fig. 2e) was comparable for all genotypes, as predicted for a TCR-dependent marker. In contrast, the signal intensity of CD25 was clearly CD28-dependent. Wildtype T cells increased CD25 MFI after CD28 costimulation while CD28^-/-^ T cells displayed lower CD25 MFI than wt cells and were unable to respond to CD28 stimulation, reflecting defective CD28 signaling. In contrast, CD25 expression was fully restored in “rescue” T cells, even after anti-CD3 stimulation alone (Fig. 2f). Finally, CD44 expression was also highly CD28-dependent. In concert with the other CD28-dependent parameters (blasting, proliferation, IL-2 and CD25), wt T cells responded to CD28 ligation with CD44 expression but CD28^-/-^ cells lacked CD44 upregulation. In contrast, the frequency of CD44^hi^CD62L^lo^ “rescue” T cells was at least comparable to fully costimulated (anti-CD3/anti-CD28) wt T cells (Fig. 2g). Thus, the miR-17~92 transgene appeared to efficiently replace CD28 function in vitro.

### Transgenic miR-17~92 enables T cell activation of CD28-deficient T cells in vivo

Next we investigated whether transgenic miR-17~92 could also substitute CD28 function in vivo. CD28^-/-^mice display a severe defect in T_FH_ development and GC formation ^24,26,28,29^. Therefore, we infected wt, CD28^-/-^ and “rescue” mice with lymphocytic choriomeningitis virus (LCMV) Armstrong to induce an acute viral infection leading to T_H_1 and T_FH_ differentiation as well as GC B cell formation. Confirming previous literature, we found severely impaired CD44 upregulation, T_FH_ differentiation and GC B cell formation in CD28^-/-^ mice compared to wt littermates (Fig. 3a-d). In contrast, all these parameters were restored in “rescue” mice (Fig. 3a-d). This is remarkable given the complexity of T_FH_ differentiation ^30^. In addition, rescued T cells not only phenotypically resembled T_FH_ cells through their expression of CXCR5, PD-1, Bcl-6 and ICOS (Fig. 3b,c) but they were functional because they induced GC B cell formation which reflects T_FH_/B cell crosstalk. Furthermore, the spleen of infected “rescue” mice, but not CD28^-/-^ mice, featured organized GCs containing GL7^+^ B cells and CD4^+^ T cells (Fig. 3e) demonstrating restoration of another hallmark defect found in CD28^-/-^ mice ^24^. Finally, we analyzed T_H_1 responses and found that transgenic miR-17~92 restored the defect in T_H_1 differentiation observed in CD28^-/-^ mice (Fig. 3f,g). We noticed that fewer CD28^-/-^ cells expressed Tbx21 compared to wt cells (Fig. 3f). However, some of those cells that did express Tbx21 co-expressed IFNγ, even in CD28^-/-^ cells. Analyzing the ratio of Tbx21^+^IFNγ^+^/Tbx21^+^ T cells confirmed that the missing costimulatory signal mainly resulted in defective Tbx21 induction rather than IFNγ production (Fig. 3h). These results suggest that the CD28^-/-^ defect acts during T cell activation, i.e. before T_H_1 differentiation. Accordingly, the miR-17~92 transgene appears to restore T cell activation signals.

**Figure 3.**
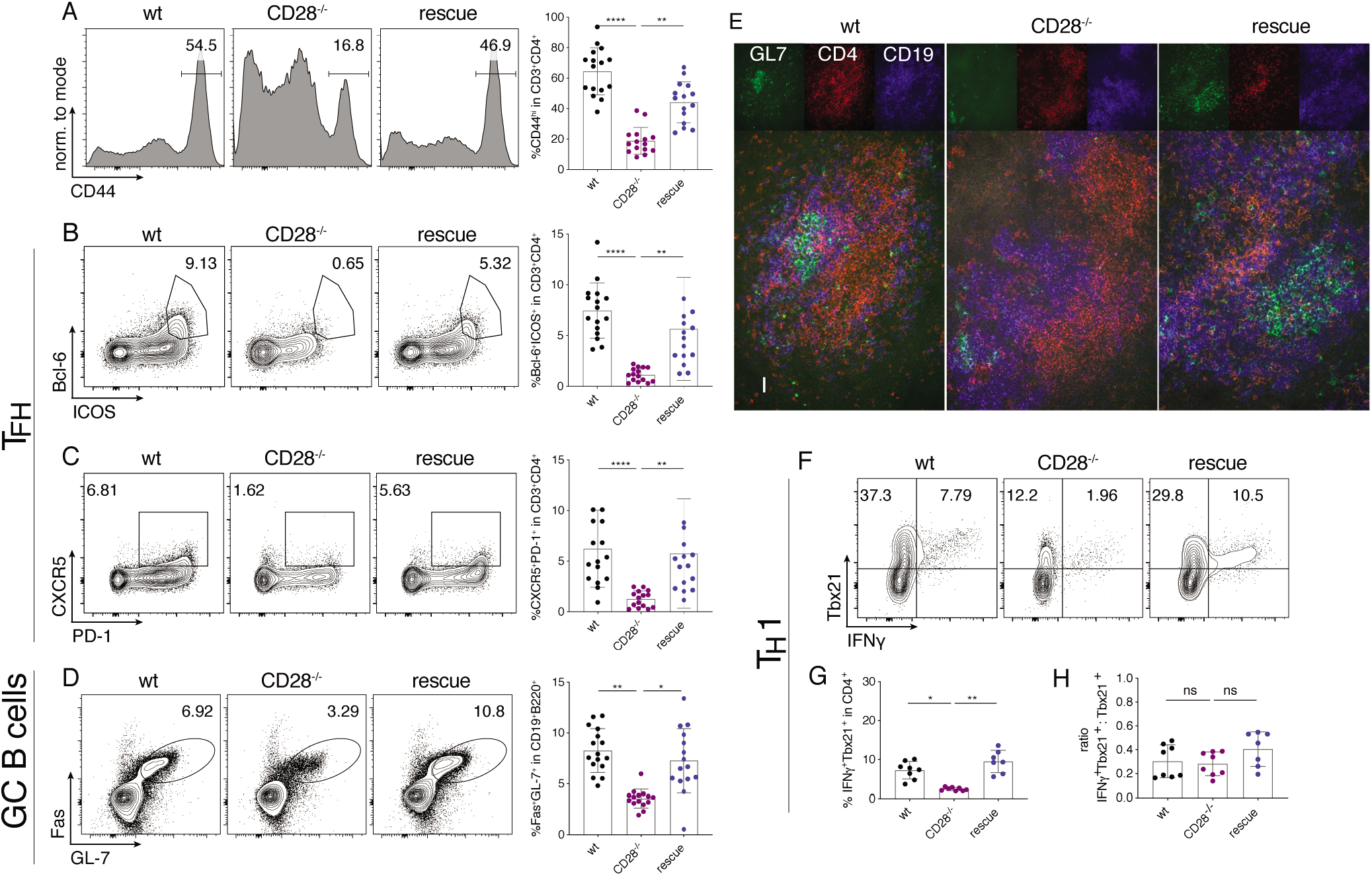
Transgenic miR-17~92 enables T cell activation of CD28-deficient T cells in vivo. 6-8 week old mice were infected with LCMV Armstrong; spleens were analyzed at d8 post infection. wt (black), CD28-/- (purple), rescue (dark blue). A-D) represent data from 4 independent experiments with 4 mice per group, pre-gating on viable CD4+CD3+ or viable CD19+B220+ cells. A) CD44 expression. B) relative number of Bcl6+ICOS+ population (TFH) C) relative number of CXCR5+PD-1+ population (TFH) D) relative number of Fas+GL7+ population (GC B cells) E) Cryosections of spleens stained for GL-7 (green), CD4 (red) and CD19 (blue). Scale bar 40μm. Splenocytes were re-stimulated with GP-64 and BFA for 4h and investigated for TH1 phenotype, pre-gated on viable CD3+CD4+ cells. Shown are 3 independent experiments with 4 biological replicates per group. F) representative contour plots of Tbx21 and IFNγ expression G) Quantification of Tbx21+IFNγ+ CD3+CD4+ cells H) ratio of Tbx21+IFNγ+ to total Tbx21 + cells. Error bars represent mean with SD, Dunn’s multiple comparison test; p values: ns=not significant, *<0.05, **<0.002, ***<0.0002, ****<0.0001

T^1792tg/tg^ T cells displayed increased proliferation and IL-2 secretion compared to wt cells (Fig. 1) and intracellular IL-2 was not only restored but even higher in “rescue” T cells than wt T cells (Fig. 2c), suggesting that the effect of miR-17~92 depended on its abundance. Therefore, we investigated the effect of a single copy of the miR-17~92 transgene in CD28^-/-^ cells. In addition, we directly compared the effect of the miR-17~92 transgene in CD28-deficient and CD28-sufficient T cells in vivo to test if CD28 triggering and the miR-17~92 transgene were additive. To this end, we infected CD28^-/-^, T^1792Δ/Δ^, wt, het rescue, rescue and T^1792tg/tg^ mice with LCMV Armstrong. “het rescue” were CD28^-/-^ that only carried one copy of the miR-17~92 Tg while “rescue” mice were the rescue mice used above (Figs. 2, 3) with two copies of the miR-17~92 transgene. The relative number of CD44^+^ T cells was lowest in CD28^-/-^ and highest in T^1792tg/tg^ mice (Suppl. Fig. 1a). We observed a similar but less pronounced effect for T_FH_, GC B cell formation and T_H_1 differentiation (Suppl. Fig. 1b-g). CD28^-/-^ and T^1792Δ/Δ^ mice exhibited a comparable defect but unlike CD44, T^1792tg/tg^ did not increase T_FH_, GC B cell formation and T_H_1 differentiation compared to wt mice. In summary, a single copy of the miR-17~92 transgene partially restored several parameters while two transgene copies further increased the rescue effect. The presence of CD28 and two transgene copies further increased IL-2 and T_H_1 formation. Thus, miR-17~92-mediated gene regulation is very sensitive to precise miR-17~92 expression in vivo.

To exclude altered viral clearance as a confounding factor we verified that wt, CD28^-/-^ and “rescue” mice normally cleared LCMV. Viral load was below detection level in plaque assays (data not shown). This is consistent with literature that LCMV clearance at this time point is CD8-mediated and CD28-independent. In summary, we found an unexpectedly complete restoration of CD28 costimulatory function and T cell activation exerted by transgenic miR-17~92 expression in vitro as well as in vivo.

### Restoration of T cell activation of CD28-deficient T cells by miR-17~92 is cell intrinsic

Although CD28’s main function is on T cells, we sought to formally test whether the miR-17~92-mediated rescue effect was cell intrinsic. We crossed MHC class II restricted CD4^+^ TCR transgenic mice specific for LCMV (SMARTA; Vα2^+^Vβ8.3^+^) to wt, CD28^-/-^ and “rescue” mice. We adoptively transferred (AT) naïve CD4^+^ T cells to CD28^-/-^ host mice followed by acute LCMV infection. Eight days post infection we isolated spleen, mesenteric and peripheral lymph nodes (LN). In all three organs the frequency and absolute number of Vα2^+^Vβ8.3^+^ CD28^-/-^ cells was strongly reduced compared to Vα2^+^Vβ8.3^+^ CD28^wt/wt^ cells. In contrast, the miR-17~92 transgene restored relative and absolute numbers of Vα2^+^Vβ8.3^+^ CD28^-/-^ T cells (Fig. 4a and Suppl. Fig. 2a, b). Furthermore, among Vα2^+^Vβ8.3^+^ T cells, fewer CD28^-/-^ cells upregulated CD44 than in wt cells, a defect that was entirely restored in rescue cells (Fig. 4b and Suppl. Fig. 2c, d). Thus, these data unequivocally demonstrate that transgenic miR-17~92 cell intrinsically compensated for CD28-deficiency. Collectively, we conclude that the non-coding RNA miR-17~92 can restore costimulatory function in CD28-deficient T cells.

**Figure 4.**
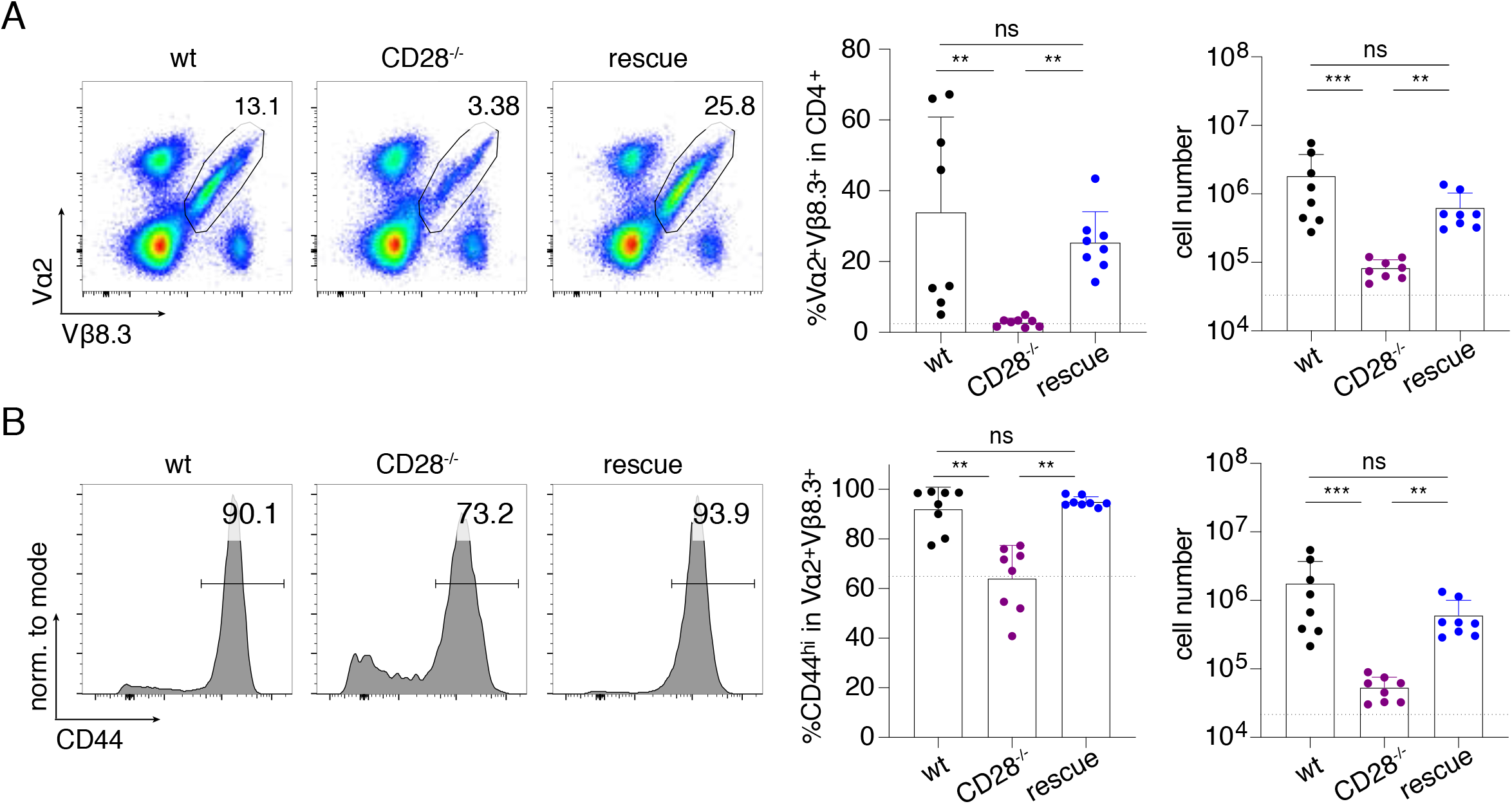
Restoration of T cell activation of CD28-deficient T cells by miR-17~92 is cell intrinsic. Adoptive transfer of naïve SMARTA+ CD4+ T cells into CD28-/- hosts, subsequent LCMV Armstrong infection and analysis of organs at d8 post infection. Donor genotypes wt (black), CD28-/- (purple), rescue (dark blue). Dotted line indicates recipient’s intrinsic Vα2+Vβ8.3+ population measured in a non-transferred control host. A) Vα2+Vβ8.3+ cells from peripheral LN pre-gated on viable, CD3+CD4+ cells B) CD44 expression in Vα2+Vβ8.3+ population from peripheral LN. 2 independent experiments, 4 recipients per group. Error bars represent mean ±SD, Dunn’s multiple comparison test, p values: ns=not significant, *<0.05, **<0.002, ***<0.0002. See also Figure S1 for spleen and mesenteric LN.

### miR-17~92 shapes the transcriptome after CD4^+^ T cell activation

To unravel the molecular mechanism underlying miR-17~92-mediated function during T cell activation we performed RNA-sequencing on naïve and in vitro activated (24h and 48h) CD4^+^ T cells from T^1792Δ/Δ^, wt and T^1792tg/tg^ mice. Principal component analysis (PCA) revealed that the transcriptomes of naïve T cells from all three genotypes were very similar (Fig. 5a, 0h). T cell activation induced major changes in gene expression (PC1, 56.7% of variance explained and PC2, 14% of variance explained) and also made the genotypes separate at 24h and even more clearly at 48h after activation (Fig. 5a; Suppl. Fig. 3a) (PC1 and PC3, 7.2% of variance explained). Since miRNAs often repress individual genes only mildly ^14^, we compared the most extreme genotypes, i.e. T^1792Δ/Δ^ to T^1792tg/tg^ to increase the power of differential gene expression analysis, at each time point. At a false discovery rate (FDR) of 1%, the number of differentially expressed genes (DEG) increased over time (830 genes up-regulated and 789 genes down-regulated at 0h, 2,493 up and 2,370 down at 24h, and 3,173 up and 3,242 down at 48h). Unsupervised hierarchical clustering of DEG 24h after activation revealed a nuanced pattern of gene clusters (Fig. 5b). As expected from the PCA (Fig. 5a), gene expression across genotypes was very similar in naïve T cells and the magnitude of expression differences increased after activation (Fig. 5b). According to their expression profile, we defined 4 different groups of genes (Fig. 5b): cluster I genes were induced over time and enhanced in T^1792tg/tg^ compared to wt but reduced or delayed in T^1792Δ/Δ^. Cluster II genes decreased with time and miR-17~92 supported their repression. Overall cluster III gene expression increased after activation but expression per time point inversely correlated with the genotype (T^1792Δ/Δ^ > T^1792tg/tg^). Thus, miR-17~92 limited initial or final maximal expression of genes in this group despite their induction with time. Finally, genes displaying the most obvious inverse correlation with the genotype were grouped in clusters IVa and IVb.

**Figure 5.**
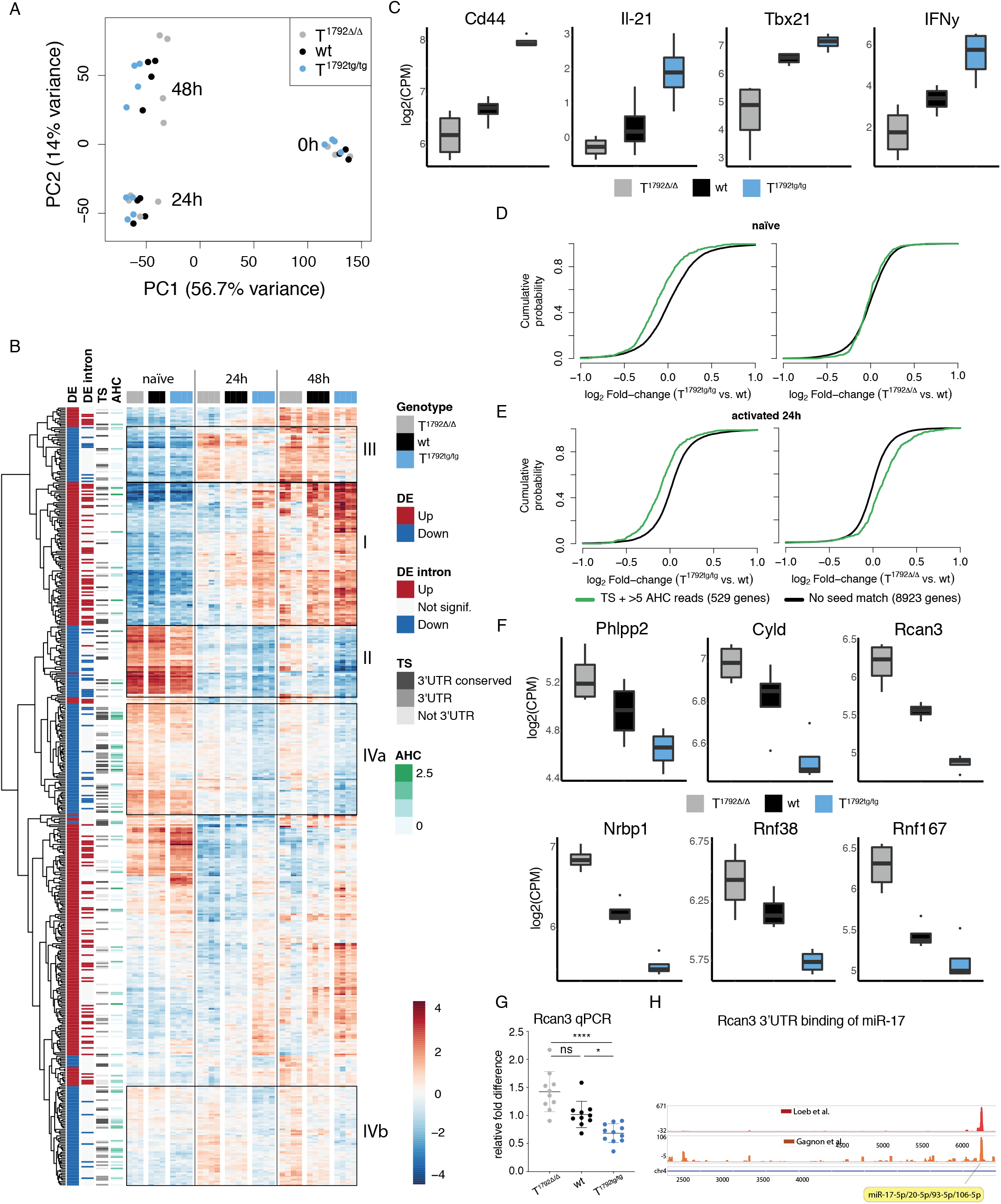
miR-17~92 shapes the transcriptome after T cell activation. Naïve CD4+ T cells from T1792Δ/Δ (grey), wt (black) and T1792tg/tg (light blue) were activated with plate-bound αCD28 and αCD3 for 0, 24h and 48h. Total RNA was extracted for bulk RNA sequencing. A) PCA (PC1 vs. PC2) based on the 25% most variable genes. B) Hierarchical clustering of the set of genes selected with abs(log2FC)>1 & adj.P.Val<0.001 in the T1792tg/tg vs. T1792Δ/Δ comparison at 24h. The heatmap displays the centered log of counts per million, with blue indicating low and red indicating high expression. Annotations: “DE” indicates the fold change direction, “DE intron” indicates if significant changes are observed in EISA analysis, “TS” indicates presence (grey) or absence (blank) of a seed match and its location and “AHC” indicates the 3’UTR signal intensity in HITS-CLIP data. Boxes I-IVb designate gene clusters. C) Examples of genes in cluster I. D, E) Genome-wide transcriptome analysis, presented as the log2 value of the gene-expression ratio for each gene versus the cumulative fraction of all log2 ratios in naïve (D) and 24h activated (E). Shown is the miR-17 seed family for T1792tg/tg vs. wt and T1792Δ/Δ vs. wt comparisons. Black curve: all genes without a seed match and ≤5 AHC reads; green: subset of genes with a seed sequence for the seed family and >5 reads in the AHC. F) Examples of genes defined as empirically validated miR-17~92 targets. log2 RNA expression level in activated CD4+ T cells, numbers correspond to FDR<0.05. G) Rcan3 mRNA expression detected by qPCR 24h after activation; shown are pooled data from three independent experiments, normalized to wt. T1792Δ/Δ (grey), wt (black), T1792tg/tg (light blue). mRNA expression normalized to 18S rRNA. Values are means ±SD, Dunn’s multiple comparison test, p values: ns=not significant, *<0.05 ****<0.0001. H) Binding sites of Argonaute 2 in 3’UTR of Rcan3 detected by HITS-CLIP; predicted miR17 binding site indicated by yellow flag. See also Figure S3 and Suppl. Table S1.

To analyze gene regulation modalities of the different clusters, we used exon-intron split analysis (EISA) to discriminate between transcriptional versus posttranscriptional regulation ^31^. In addition, we employed computational target gene predictions from the Targetscan (“TS”) database ^32^ and used a dataset of biochemically detected direct miRNA:mRNA interactions in T cells defined by Argonaute 2 high-throughput sequencing of RNA isolated by crosslinking immunoprecipitation (“AHC”) ^33^. For each gene we quantified the read coverage on TS seed matches for each of the miR-17~92 cluster seed families. These predictors of transcriptional (“DE intron”) and posttranscriptional regulation (“TS”, “AHC”), annotated on the heatmap (Fig. 5b) revealed genes whose expression increased over time and positively correlated with the miR-17~92 genotype. They were enriched for transcriptional regulation and displayed few TS sites or AHC reads (Fig. 5b, box I). Thus, cluster I was mainly induced by increased gene transcription and miR-17~92 promoted this transcriptional activity. Examples include *Cd44, IL-21, Tbx21* and *IFNγ* (Fig. 5c). In contrast, genes from clusters IVa and IVb (Fig. 5b, boxes IVa, IVb) displayed a consistent and colinear reduction of expression levels across genotypes that inversely correlated with miR-17~92 dosage (T^1792Δ/Δ^ > wt > T^1792tg/tg^) at 24h and 48h. Importantly, these clusters contained few transcriptionally regulated genes but were enriched for TS sites (p-value=0.001037; Fisher test) and experimentally determined AHC reads (p-value=0.003598; Fisher test) (Fig. 5b). Thus, these clusters were mainly regulated posttranscriptionally and likely constituted direct miR-17~92 target genes.

To further characterize direct miR-17~92 target genes we focused on naïve T cells and the 24h time point since indirect effects likely increased after this time point. To visualize the effect of individual miR-17~92 cluster miRNAs on their target genes we compared expression of genes identified by TS and with >5 AHC reads for each miRNA seed family to all genes without any seed match for that family. As illustrated by the miR-17 seed family, the miR-17~92 transgene repressed miR-17 target genes in naïve T cells (Fig. 5d, left panel) but absence of miR-17~92 had no effect on the expression of miR-17 target genes (Fig. 5d, right panel). In contrast, after T cell activation miR-17 target genes were repressed in T^1792tg/tg^ T cells (Fig. 5e, left panel) and derepressed in T^1792Δ/Δ^ T cells (Fig. 5e, right panel). Similar effects were observed for all seed families although to a lesser extent for the miR-18 seed family (Suppl. Fig. 3b, c). We defined genes as empirically validated miR-17~92 targets if they fulfilled the following criteria: i) significant derepression in T^1792Δ/Δ^ vs wt and significant repression in T^1792tg/tg^ vs wt at 24h ii) predicted TS match iii) >5 AHC reads and iv) posttranscriptional regulation based on EISA. Applying these criteria across the 4 seed families from miR-17~92 defined a set of 68 empirically supported direct miR-17~92 target genes (Suppl. Table 1). These genes included previously described miR-17~92 targets validated in T cells, such as *Phlpp2* ^23^ and *Cyld,* validated in B cells ^34^ (Fig. 5f). In addition, this approach identified many less studied genes (e.g. *Rcan3, Nrbp1, Rnf38 and Rnf167*) as miR-17~92 targets in T cells (Fig. 5f). Notably, several of the target genes negatively regulate pathways important for T cell activation. For instance *Phlpp2* is a PI3K inhibitor ^23^, *Cyld* is a NF-κB inhibitor ^35^ and *Rcan3* a putative calcineurin inhibitor ^36^ (Fig. 5f). Since miR-17~92 has not been reported to regulate calcineurin signaling in T cells, we validated the *Rcan3* RNA-Seq data by qPCR on independent biologic replicates (Fig. 5g). Moreover, its 3’UTR contains a conserved miR-17-5p 8mer binding site and AHC confirmed a single discrete peak at this site, evidence for a direct interaction of a miRNA with the *Rcan3* 3’UTR in primary T cells (Fig. 5h and ^15,33^). Together, these data empirically validate *Rcan3* as a miR-17 target in T cells.

Thus, using a combination of experimentally validated differential gene expression (T^1792Δ/Δ^, wt and T^1792tg/tg^ T cells), evidence of posttranscriptional gene regulation and biochemical detection of miRNA binding we defined a high confidence set of miR-17~92 target genes in T cells. miR-17~92 became functionally relevant after T cell activation and shaped the transcriptome in intricate ways. Our data demonstrates that miR-17~92 can promote and inhibit gene expression. miR-17~92-mediated transcriptional consequences are most likely indirectly mediated through posttranscriptional gene regulation. In addition, direct inhibition can enhance gene silencing or dampen expression of induced genes. Thus, miR-17~92-mediated gene repression is very powerful to shape the T cell transcriptome during T cell activation.

### miR-17~92 promotes the calcineurin/NFAT pathway in CD4^+^ T cells

To identify which molecular pathways were regulated by miR-17~92, we analyzed curated gene sets enriched for differentially expressed genes between T^1792tg/tg^ and T^1792Δ/Δ^. At 24h, the gene sets with the highest statistical significance and largest average fold change were related to cytokines, inflammation and T cell differentiation (Suppl. Fig. 4a) while at 48h many metabolic pathways were altered (Suppl. Fig. 4b). Next, we performed enrichment analysis on DoRothEA regulons ^37^ to identify TF activity that could explain the regulated pathways. Regulons were defined using any existing interaction with a particular TF. At 24h, the five most significantly enriched TF regulons with the highest fold change contained two NFAT members (NFATC2, NFATC3) as well as RELA, NF-κB1 and GATA3 (Fig. 6a). Since NFAT TFs are important for T cell activation and differentiation (including T_FH_ differentiation ^38^) but miR-17~92 is not known to promote NFAT activity in T cells, we focused on the calcineurin/NFAT axis. Genes belonging to regulons NFATC2 and NFATC3 include many T cell lineage-defining TFs, cytokines and cytokine receptors and most of them – including *IL-21, Tbx21* and *IFNγ* (Fig. 5c) - were highly regulated by miR-17~92 (Fig. 6b). This confirmed that miR-17~92 constitutes a central regulator of T cell activation and suggested that transgenic miR-17~92 enhanced canonical pathways that resulted in functional substitution of CD28 for differentiation of T_FH_ and T_H_1 in vivo (Fig. 3). Since miR-17~92 promoted expression of signature TF and cytokines defining multiple T cell subsets, we tested if miR-17~92 more generally could replace CD28. To this end we differentiated T_H_1, T_H_17 and iTreg cells in vitro. We confirmed that also in this setup transgenic miR-17~92 was sufficient to compensate for the absence of CD28 and functionally corrected the defects of CD28^-/-^T cells to differentiate into all 3 subsets (Suppl. Fig. 5).

**Figure 6.**
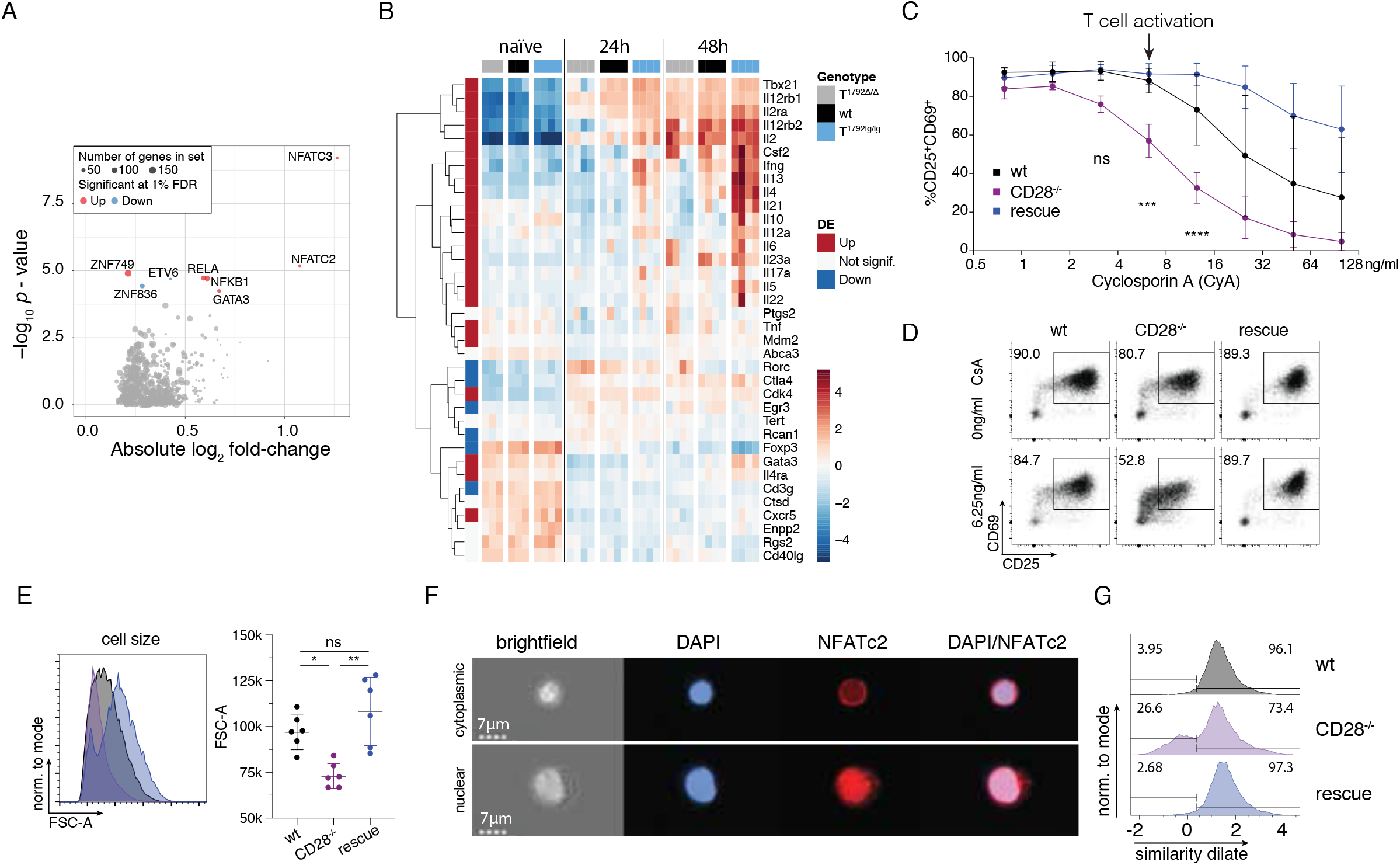
miR-17~92 promotes the calcineurin/NFAT pathway in CD4+ T cells. A, B) Dataset from Figure 5. Naïve CD4+ T cells from T1792Δ/Δ (grey), wt (black) and T1792tg/tg (light blue) were activated with plate-bound αCD28 and αCD3 for 0, 24h and 48h. Total RNA was extracted for bulk RNA sequencing. A) Volcano plot showing the absolute log2 fold change and −log10 p value from regulon analysis. A threshold of 1% FDR was applied. Dot size indicates number of genes within each regulon and colors indicate fold change direction. B) Heatmap of genes under NFATC2_D and NFATC3_D regulons plus several known activated genes in CD4+ T cells (Il10, Il12a, Il6, Rorc, Il23a). Hierarchical clustering was applied on genes. Centered log of counts per million is displayed, with blue indicating low and red indicating high expression. C) CD4+ wt (black), CD28-/- (purple) and rescue (dark blue) cells were activated for 48h in the presence of increasing concentrations of cyclosporin A (CyA) as indicated and stained for CD25 and CD69. Shown are 2 independent experiments, error bars represent means ±SD. Tukey’s multiple comparison, p values: **<0.002, ****<0.0001 refer to the difference between CD28-/- and wt. D) representative FACS plots of CD25/CD69 expression in viable CD4+ T cells activated for 48h with no or 6.25ng/ml CyA. E) influence of 6.25ng/ml CyA on blasting (FSC-A of the lymphocyte gate) of viable CD4+ cells. F) Imagestream analysis of CD4+ T cells, activated for 48h in presence of 6.25ng/ml CyA; DAPI (blue) and NFATc2 (red) staining. Examples for cytoplasmic (top) and nuclear (bottom) NFATc2 in a CD28-/- sample. G) histograms of the similarity dilate indicative of the co-localization of NFATc2 and DAPI signals; gates indicate the nuclear (high similarity dilate) and the cytoplasmic population (low similarity dilate). See also Figures S4 – S6.

While miR-17~92 predominantly promoted inflammatory pathways at 24h, at 48h several metabolic pathways were regulated by miR-17~92 (Suppl. Fig. 4a, b). Thus, we hypothesized that initial transcriptional changes could lead to altered metabolism. It was recently shown that cell cycle entry of quiescent T cells was controlled by store operated Ca^2+^ entry (SOCE) and calcineurin/NFAT through control of glycolysis and oxidative phosphorylation ^39^. Therefore, we analyzed T cell metabolism in naïve and activated T^1792Δ/Δ^, wt and T^1792tg/tg^ T cells. In line with the trancriptome results (Fig. 5), metabolic flux analysis demonstrated comparable glycolytic and respiratory activity of naïve T cells in T^1792Δ/Δ^, wt and T^1792tg/tg^ T cells (Suppl. Fig. 6a, b). In contrast, 48h after activation, glycolytic and respiratory activity positively correlated with the miR-17~92 genotype (Suppl. Fig. 6c, d). Furthermore, genes associated with the TCA cycle and respiratory electron transport positively correlated with miR-17~92 at 48h but not before, supporting the notion that miR-17~92 indirectly promoted this metabolic activity (Suppl. Fig. 6e).

Since transcriptome analysis identified calcineurin/NFAT as a key pathway promoted by miR-17~92 (Fig. 6a, b), we designed experiments to functionally validate this increased activity. We activated wt, CD28^-/-^ and “rescue” T cells in the presence of cyclosporin A (CyA), a direct pharmacological calcineurin inhibitor (CNI) and quantified CD69/CD25 expression. Compared to wt cells, CD28^-/-^ cells were 4-fold more sensitive to CyA (Fig. 6c). At a CyA concentration that did not affect wt cells (arrow in Fig. 6c), T cell activation of CD28^-/-^ T cells was clearly inhibited (Fig. 6c,d). Thus, compared to wt cells CD28^-/-^ T cells are hypersensitive to CyA. In contrast, CD28^-/-^ T cells with forced miR-17~92 expression were normally activated (Fig. 6c, d). In fact, at higher CyA concentrations the “rescue” T cells were even more resistant to CyA inhibition than wt T cells (Fig. 6c). These findings were further corroborated by the analysis of cell size which revealed that in the presence of low dose CyA the blasting defect of CD28^-/-^ T cells was compensated for by transgenic miR-17~92 (Fig. 6e). Finally, we assessed nuclear translocation of activated NFATC2 as a direct readout of calcineurin activity. ImageStream analysis of T cells activated in the presence of 6.25ng/ml CyA showed T cells with cytoplasmic (top row) or nuclear (lower row) NFATC2 (Fig. 6f). Quantification of this data demonstrated that the presence of a low CyA concentration reduced nuclear NFATC2 translocation in CD28^-/-^ T cells but was restored in “rescue” cells (Fig. 6g). Thus, the miR-17~92 transgene replaced CD28-enhanced calcineurin signals for blasting, CD69 upregulation and nuclear translocation of NFATC2.

### miR-17~92 is necessary for repression of a subset of genes during T cell activation

Finally, we examined whether miR-17~92 was physiologically required to shape the molecular program triggered by CD28 engagement. We repeated transcriptome analysis with RNA-seq on naïve and activated (24h) CD4^+^ T cells from T^1792Δ/Δ^, wt and T^1792tg/tg^ mice but added CD28^-/-^ and “rescue” T cells. The correlation of miR-17~92 target gene expression levels with the first RNA-seq experiment (Fig. 5) was very high (Suppl. Fig. 7a). A PCA showed that the transcriptomes of naïve T cells of all 5 genotypes were closely related (Fig. 7a). However, after T cell activation (PC1, 67.6% of variance) the 5 genotypes formed 5 distinct groups that were separated on PC2 (5.6% of variance). Notably, the “rescue” samples were located between CD28^-/-^ and T^1792Δ/Δ^, wt and T^1792tg/tg^ T cells (Fig. 7a). This implies that transgenic miR-17~92 partially restored the genetic networks dysregulated by CD28-deficiency. Thus, the PCA analysis supported the notion that the phenotypic rescue (Figs. 2-4) was a consequence of a partially restored transcriptome rather than a transgene artifact. Moreover, compared to all genes without a miR-17~92 seed match the miR-17~92 target gene set defined earlier (Suppl. Table 1) was overall derepressed in CD28^-/-^ T cells compared to wt cells (Fig. 7b and Suppl. Fig. 7b). It is important to note that these cells have an untouched miR-17~92 locus and therefore should repress miR-17~92 target genes like wt cells. However, the absence of CD28 resulted in increased expression of those genes. This result demonstrated that CD28-dependent effects normally rely on miR-17~92 to repress dozens of genes during T cell activation. Remarkably, expression of target genes was corrected in “rescue” cells (Fig. 7c and Suppl. Fig. 7b). Thus, transgenic miR-17~92 repressed the physiologic targets that were derepressed in CD28^-/-^ T cells.

**Figure 7.**
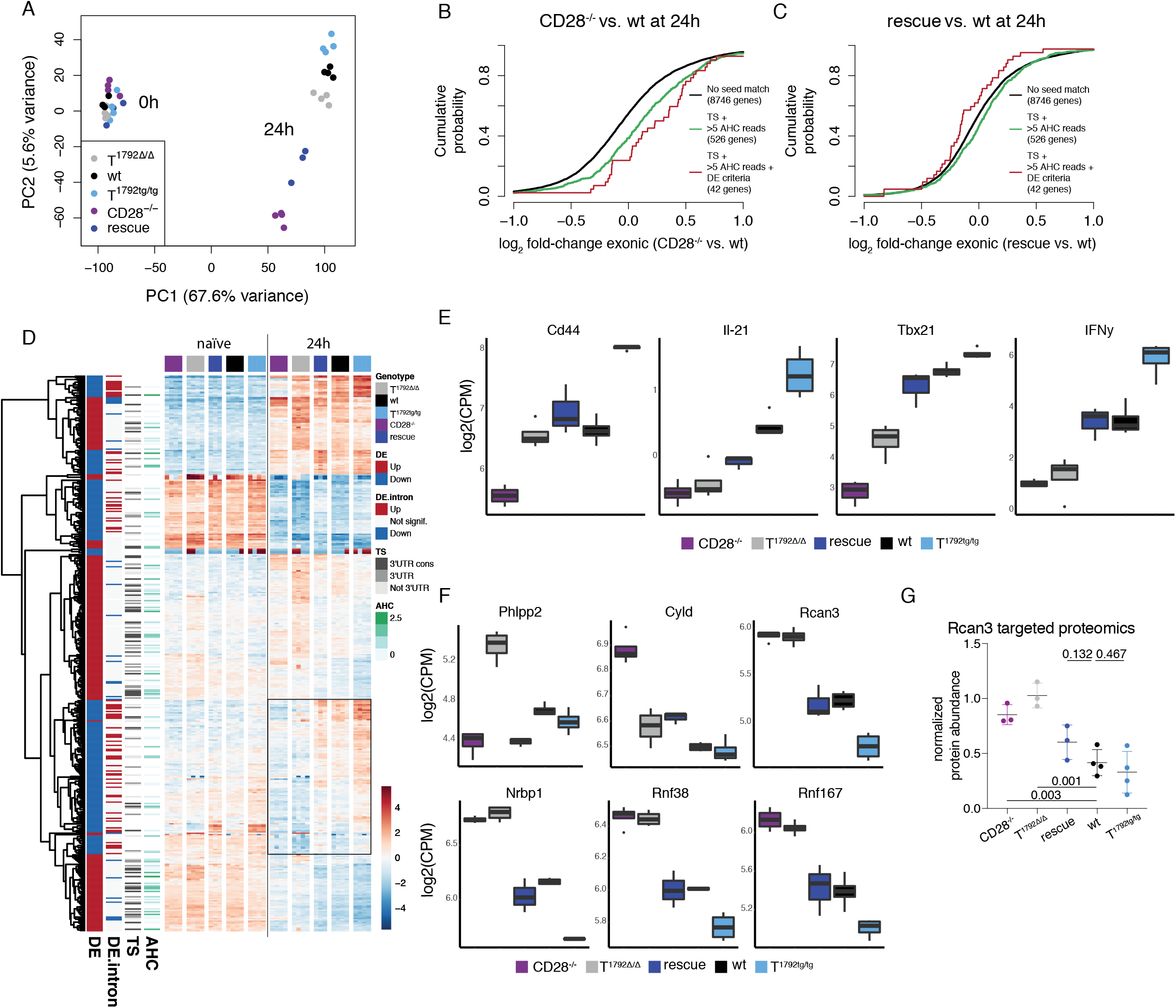
miR-17~92 is necessary for repression of a subset of genes during T cell activation. CD4+ T cells from T1792Δ/Δ (grey), wt (black), T1792tg/tg (light blue), CD28-/- (purple) and rescue (dark blue) mice were activated for 24h. Total RNA was extracted for sequencing. A) PCA (PC1 vs. PC2) based on the 25% most variable genes. B) Genome-wide transcriptome analysis, presented as the log2 value of the gene-expression ratio for each gene versus the cumulative fraction of all log2 ratios. Shown are the contrasts between activated samples separated by the miR-17 seed family for the comparison CD28-/- vs. wt (B) and rescue vs. wt (C). Black curve: genes without a seed match, ≤5 AHC reads, and no differential expression in the second RNA sequencing. Green: genes with a conserved binding site for the miR-17 seed family (TS) and >5 reads in the AHC, red: genes with a conserved binding site for the miR-17 seed family (TS), >5 reads in the AHC and differential expression in the second RNA sequencing data set. D) Hierarchical clustering of the set of genes selected with abs(log2F-C)>1 & adj.P.Val<0.001 in the T1792tg/tg vs. T1792Δ/Δ comparison at 24h. The heatmap displays the centered log of counts per million, with blue indicating low and red indicating high expression. E) Examples of transcripts that were restored in “rescue” compared to CD28^-/-^ T cells, i.e. contained in the box of Fig. 7d. F) Examples of direct miR-17~92 targets. E, F: log2 mRNA expression in activated CD4+ T cells; numbers correspond to FDR<0.05. G) Rcan3 protein abundance measured by targeted proteomics 24h after activation. CD28-/- (purple), T1792Δ/Δ (grey), rescue (dark blue), and T1792tg/tg (light blue) are compared to wt (black). Proteins were isolated from the same cells for targeted proteomics of Rcan3 and total RNA sequencing. Numbers indicate p values from t-test. See also Figure S7.

However, the molecular rescue effect on the entire transcriptome was incomplete (Fig. 7a). In order to determine which transcripts were restored and whether they were regulated transcriptionally or posttranscriptionally, we performed unsupervised hierarchical clustering of the same genes as shown in Fig. 5b but used the gene expression values from the 2^nd^ dataset with all 5 genotypes. At 24h, one gene cluster stood out in which “rescue” T cells were more similar to wt and T^1792tg/tg^ cells than CD28^-/-^ and T^1792Δ/Δ^ cells (Fig. 7d, box). Genes in this cluster contained several NFAT-dependent transcripts including *IL-4, IL-12a, IL12rb2, IL-21* and *IFNγ* (Suppl. Table II, Fig. 6b). Importantly, *Cd44, IL-21, Tbx-21 and IFNγ* transcripts, also contained in this cluster, were reduced in CD28^-/-^ compared to wt cells (Fig. 7e). In contrast, transgenic miR-17~92 partially or completely restored these transcripts in “rescue” cells to wt levels. Furthermore, T^1792tg/tg^ T cells expressed supraphysiologic mRNA levels (Fig. 7e). Thus, the phenotypic rescue of CD44 and IFNγ in CD28^-/-^ T cells (Fig. 2-4) can at least partially be attributed to increased expression of these genes driven by the miR-17~92 transgene in “rescue” cells.

Conversely, as illustrated above (Fig. 7b) many direct targets were not only derepressed in T^1792Δ/Δ^ but also in CD28^-/-^ cells. Several genes (e.g. *Rcan3, Nrbp1, Rnf38 and Rnf167)* were similarly derepressed in CD28^-/-^ and T^1792Δ/Δ^ cells and expression was restored in “rescue” cells to levels similar to wt and even further repressed in T^1792tg/tg^ cells (Fig. 7f). Others, such as *Phlpp2* and *Cyld* were clearly regulated by miR-17~92 but expression differed in CD28^-/-^ and T^1792Δ/Δ^ cells (Fig. 7f). For one of these targets, *Rcan3*, we confirmed this expression pattern using targeted proteomics for absolute quantification (Fig. 7g). This suggests that these targets are sensitive to regulation by miR-17~92 and their derepression in CD28^-/-^ T cells is largely attributed to reduced repression by miR-17~92. Collectively, the data demonstrates that miR-17~92 is necessary for CD28-induced gene repression.

## Discussion

T cell activation depends on TCR and CD28 engagement to trigger complex molecular mechanisms including activation of the PI3K, NF-κB and calcineurin/NFAT pathways. Intense research in the past decades uncovered many molecules that transmit TCR and CD28 signals ^2^. Most studies focused on proteins as signaling intermediates but we and others previously reported that combined engagement of TCR and CD28 also alters expression of non-coding RNAs. Most miRNAs are downregulated after T cell activation but a few, including miR-17~92, remain relatively constant or are slightly induced ^20,40^. This suggests that miR-17~92 might be functionally relevant during T cell activation. We previously found that miR-17~92 was induced after combined stimulation of TCR and CD28 in vitro but not after TCR stimulation alone ^20^. These findings were confirmed and extended to the natural ligands CD80/CD86. Heterozygosity or absence of CD80/CD86 on B cells resulted in a dose-dependent reduction in miR-17 induction in co-cultured T cells while deficiency or blockade of CTLA-4 resulted in increased miR-17 expression ^41^. Furthermore, stimulating T cells with antibodies directed to CD3 is sufficient to induce proliferation in T^1792tg/tg^ T cells ^21^. Collectively, these studies suggest that T cell activation, CD28 and miR-17~92 are intimately linked. However, their relation is poorly understood. Here, we tested whether transgenic miR-17~92 was sufficient to substitute for the absence of CD28 and whether miR-17~92 was required for CD28-mediated T cell activation. We found an unexpectedly potent rescue effect both in vitro and in vivo. Many defects of CD28^-/-^ T cells were functionally compensated for by the miR-17~92 transgene. This is notable for two reasons. First, miR-17~92 is a non-coding RNA and second, miRNAs are negative regulators. Thus, overexpression of an inhibitory RNA was sufficient to enable T cell activation in the absence of CD28. However, the miR-17~92 transgene did not equally restore all investigated parameters. For instance, the transcriptome was incompletely restored while i.c. IL-2 and the relative number of CD44^+^ cells were even higher in T^1792tg/tg^ than “ “rescue” T cells. Moreover, although *Phlpp2* was clearly regulated by miR-17~92, it was not derepressed in CD28^-/-^ T cells. Thus, CD28 must control pathways that are independent of miR-17~92 and not all molecular programs are equally sensitive to miR-17~92 regulation. Nevertheless, we observed a graded rescue of some parameters with a single or two copies of the miR-17~92 transgene. This data is consistent with recent findings that sensing of costimulatory ligands in vivo is analog, not digital. Commitment to T_FH_ differentiation appears to require a higher level of CD28 engagement than commitment to proliferation or CD62L downregulation ^41^. The tight connection between CD28 and miR-17~92 ^REFS 20,41^ together with the data described here might explain why T_FH_ are particularly sensitive to miR-17~ 92 ^22,23^ and opens the possibility that miRNAs of the miR-17~92 cluster could be leveraged to tune T cell activation/costimulation.

To analyze the molecular mechanism by which miR-17~92 enabled T cell activation in the absence of CD28 we took advantage of a multi-layer approach. RNA-Seq analysis of T cells with miR-17~92 loss- or gain of function and sampled over a time course revealed that miR-17~92 mainly influenced the transcriptome after T cell activation, a finding consistent with phenotypic analysis of ex vivo characterized naïve T cells. There is an emerging notion that evidence of miRNA binding combined with differential gene expression in primary cells constitutes a precise approach to empirically define miRNA:target relationships ^16,33^. We refined this approach by combining EISA with AHC to discriminate whether differentially expressed gene clusters were primarily regulated transcriptionally or posttranscriptionally. This analysis provides a highly granular view of miRNA-mediated gene regulation. Specifically, pathway and regulon enrichment analysis revealed that miR-17~92 enhanced the calcineurin/NFAT pathway. Consistent with this, a gene cluster enriched for genes that positively correlated with miR-17~92 was mainly transcriptionally regulated and contained many genes known to be regulated by NFAT TF including genes for various CD4^+^ T cell subsets such as T_FH_ (*IL-21*) and T_H_1 (*Tbx-21 and IFNγ*). Experiments with pharmacologic CNIs confirmed that miR-17~92 functionally promoted the calcineurin/NFAT pathway. Importantly, transgenic miR-17~92 partially restored the molecular program that was defective in CD28^-/-^ T cells. In particular, expression of key NFAT-regulated genes found to be driven by miR-17~92 in CD28-sufficient T cells were restored by the transgene in CD28-deficient T cells. Although CD28 is critical for early T cell priming, CD28 costimulation continues to be important for days ^4^. Our data shows that in the first 24h miR-17~92 has important functions by promoting T cell activation and expression of TF, cytokines and cytokine receptors required for differentiation into multiple T cell subsets. We and others previously demonstrated that miR-17~92 also supports T cell differentiation and/or subset specific functions at later time points, e.g. by repressing lineage inappropriate TF and facilitating PI3K signaling ^22,23^. We propose that miR-17~92 acts as a downstream mediator or amplifier of the molecular program triggered by combined TCR/CD28 engagement. However, how CD28/T cell activation is linked molecularly to miR-17~92 expression remains poorly understood. Although the presence of myc binding sites is suggestive for transcriptional regulation ^42^, miRNA biogenesis can be controlled by extracellular cues ^43^ and in embryonic stem cells maturation of miR-17~92 into mature miRNAs is also regulated posttranscriptionally ^44^. Future studies are needed to unravel how CD28 costimulation/T cell activation regulate miR-17~92 expression. Nevertheless, our data demonstrates that miR-17~92, induced by the combined triggering of TCR and CD28, is required to mediate gene repression that indirectly promotes gene expression. Thus, miR-17~92 is part of a feed forward loop.

How can a microRNA cluster that negatively regulates genes promote gene expression and bypass the requirement for CD28 costimulation? To shed light on this question we first defined miR-17~92 target genes in T cells using miR-17~92 gain- and loss of function as well as AHC. Next we demonstrated that these same genes were elevated in stimulated CD28^-/-^ T cells. We conclude that during T cell activation miR-17~92 is required to repress this set of genes. Previous work demonstrated that miR-17~92 promotes the PI3K, NF-κB and mTOR pathways. Here we demonstrate that it also enhances the calcineurin/NFAT pathway. We noticed that the identified, empirically validated miR-17~92 target gene network contains multiple negative regulators of the PI3K (*Phlpp2*), NF-κB (*Cyld*) and NFAT (*Cyld, Rcan3*) pathways. Furthermore, the list contains poorly characterized known or suspected tumor suppressors, genes related to cell growth, proliferation, migration and cancer (e.g. *Nkiras1, Nrbp1, Tgif1, Ugp2, Wdr26, Zbtb4*) as well as known or suspected E3 ligases (*Ankib1, Dcaf8, Rnf38* and *Rnf167*). Since miR-17~92 promotes proliferation and can be oncogenic it is suggestive that repression of at least some of these genes contributes to proliferation after T cell activation.

The importance of removing negative regulators to enable T cell activation is illustrated by limited experimental evidence that gene deletion can partially rescue CD28-deficiency. For instance, genetic deletion of Casitas B lymphoma-b protein (*Cbl-b*) is sufficient to restore IL-2 production and proliferation in CD28^-/-^ T cells. However, neither IFNγ, IL-4 or ICOS expression nor GC formation were restored ^45,46^. Thus, while loss of Cbl-b uncoupled T cells from the strict requirement for CD28 costimulation, it could not restore all aspects of CD28-mediated costimulation. Similarly, genetic ablation of PTEN or TRAF6 restored proliferation and IL-2 production in CD28^-/-^ CD4^+^ T cells ^47,48^. In addition, peripheral Cyld^-/-^ T cells display increased proliferation, IFNγ and IL-2 production upon activation. Thus, Cyld-deficient T cells are hyperresponsive to stimulation and therefore Cyld is another important restrainer of T cell activation ^35^. Because of these genetic examples it was noted before that the default consequence of TCR triggering is preventing T cell activation and induction of anergy ^49^. As a group of genes, negative regulators impose a requirement for a costimulatory signal to actively remove these brakes to allow a productive T cell response. CD28 costimulation therefore serves a dual purpose by enhancing TCR signaling and simultaneously overcoming T cell repression ^12,48,49^. Interestingly, *Cbl-b, PTEN* and *Cyld* are all directly targeted by miR-17~92. Importantly, during T cell activation the inhibitors are not completely eliminated as is the case in the genetic deletion systems. Rather, TRAF6, PTEN and Rcan3 expression decreases but these genes remain expressed at substantial levels and Cbl-b expression even increases (induced by NFAT) ^47^. Thus, although experimental deletion can demonstrate that a given gene has the potential to act as a powerful negative regulator, its physiologic function is more delicate to uncover and complete deletion likely overestimates the contribution of the gene under investigation. To date no single gene or pathway can explain the function of CD28 costimulation. More likely, multiple negative regulators need to be removed and multiple pathways need to be induced. Our results illustrate that miR-17~92 can exert complex regulation of the T cell transcriptome through direct and indirect gene regulation. Mammalian gene expression is rarely digital and controlled by multiple layers of molecular mechanisms. By combining transcriptional with posttranscriptional gene regulation, animals can achieve complex gene expression with relatively simple promoters ^14^. It is conceivable that precise control of genetic programs is particularly important for an exponential process like clonal T cell expansion. Even minor changes in setting the T cell activation threshold or the rate of proliferation or survival could result in major consequences for the host organism. Since miRNAs are important sculptors of the transcriptome they are ideally suited to mediate controlled release from multiple restraining proteins preventing T cell activation ^14^. Therefore, genes like *Rcan3, Nrbp1, Rnf38* and *Rnf167* are of particular interest for further investigation since they are equally derepressed in CD28^-/-^ and T^1792Δ/Δ^ CD4^+^ T cells but very sensitive to regulation by miR-17~92 since their expression in “rescue” cells is similar to wt cells. It will be interesting to determine if their repression contributes to bypassing CD28 in “rescue” T cells. Other miR-17~92-regulated genes such as *Cyld* are also of interest but miR-17~92-independent mechanisms must be operational since its expression is not affected as much in T^1792Δ/Δ^ as in CD28^-/-^ CD4^+^ T cells. Clearly, additional studies are needed to decipher the relative contributions of each individual or combinations of target genes in primary T cells and oncogenesis because miRNA regulation is highly cell type and context dependent ^16^. Nevertheless, our findings greatly expand the list of high confidence miR-17~92 targets with a suspected function in T cells.

In summary, we propose a model in which miR-17~92 constitutes an important mediator of T cell activation/CD28 costimulation by serving as a gatekeeper of a network of restrainers controlling at least 3 central pathways for T cell activation and function. In this model, non-coding RNA-mediated repression of inhibitors supports transcriptional induction of genes necessary for further T cell differentiation and function (e.g. CD44, IL-21, Tbx21, IFNγ). Importantly, we not only show that transgenic miR-17~92 is largely sufficient to substitute for CD28-deficiency but that miR-17~92 is physiologically required to shape the transcriptome after CD28 costimulation and that miR-17~92 overexpression in CD28-sufficient T cells results in “super-costimulation”. In support of the proposed model, constitutive transgenic miR-17~92 expression leads to a lupus-like systemic autoimmune syndrome ^21^, not unlike the ones observed in mice deficient in *Cbl-b, PTEN, TRAF6* or *Cyld* ^35,45,46,50,51^. Our results support the consideration of miR-17~92 or its target genes for pharmacologic intervention or for its integration into engineered cellular therapies.

## Data and materials availability

High-throughput RNA sequencing data will be publically available at GEO accession GSE140568 once published in a peer reviewed journal.

## Acknowledgements

We would like to thank the current and former lab members for scientific discussions; Daniel Pinschewer and Weldy Bonilla Pinschewer for advice on LCMV, reagents (GP61 peptide) and scientific discussions; Annaïse Jauch for providing LCMV Armstrong; Mihaela Zavolan and Ed Palmer for deep scientific discussions and critical comments on the manuscript; members of the NCCR RNA & Disease for discussions; the University of Basel, the Basel University Hospital and Department of Biomedicine (DBM) for institutional support; the team from the DBM animal facility for animal husbandry; the DBM flow cytometry core team; Robert Ivanek from the DBM bioinformatics core; Philippe Demougin from the D-BSSE/DBM Genomics Facility Basel; Alexander Schmidt from the University of Basel Biozentrum Proteomics Core Facility for support and Marco Amsler, Caroline Schwenzel, Hélène Rossez, Giuseppina Capoferri and Corinne Engdahl-Démollière for technical assistance. Calculations were performed at sciCORE (http://scicore.unibas.ch/) scientific computing center at University of Basel. This work was supported by grants of the Swiss National Science Foundation (SNSF Professorship PP00P3_144860 to LTJ) and the National Institute Of Allergy And Infectious Diseases of the National Institutes of Health, USA, under Award Number R56/R01AI106923 (to LTJ). The content of this study is solely the responsibility of the authors and does not necessarily represent the official views of the National Institutes of Health.

## Author contributions

M.D. performed and analyzed most experiments; G.B. and C.H. helped to design and interpret metabolic experiments; R.M. provided scientific input, key experimental and infrastructure support; M.K. provided scientific input and experimental support; R.K. generated HITS-CLIP data; D.S. and J.R. performed bioinformatic analysis with the help of J.G.; K.M.A. provided key scientific input and access to HITS-CLIP data; M.D. and L.T.J. designed the experiments, interpreted the data and discussed results; L.T.J. designed the study, obtained funding, supervised all experiments and wrote the manuscript (MS) with the help of M.D.; all authors approved the final MS.

## Animals

### Mice

All animal work was performed in accordance with the federal and cantonal laws of Switzerland. Protocols were approved by the Animal Research Commission of the Canton of Basel-Stadt, Switzerland. Most of the mouse lines were imported from the JAX laboratory as indicated in the key resources table. T^1792Δ/δ^ mice were imported from the laboratory of Dr. Bluestone (UCSF). Rescue SM^+^ mice were obtained by crossing the rescue strain to other transgene lines in house so that their precise genetic origin is difficult to determine. As for the transfer cells, we used SM CD45.1^+^ wt cells imported from SWIMR. All other mouse lines were crossed to CD45.2 expression. Cre negative littermates (from T^1792tg/tg^ or T^1792Δ/δ^) were used as wt controls, and cre negative littermates from the rescue strain were used as CD28ko. 6-8 week old females and males were used for all experiments.

## METHOD DETAILS

### Organ isolation

Organs were obtained after CO_2_ euthanization and kept on ice until processing. Mesenteric lymph nodes (LN), peripheral LN (inguinal, axillary, brachial, six cervical) and spleen were taken for most of the experiments. Spleens were injected with 0.5ml ACK lysis buffer (0.155M NH4Cl, 200μl 0.5M EDTA pH=8.0, 0.012M NaHCO_3_ pH 7.2) for erythrolysis before processing. The organs were meshed with 0.4μm filters to obtain single cell suspensions which were then washed with FACS buffer (2% heat-inactivated FCS in PBS, for stainings add 0.02% NaN3). Cells were centrifuged at 4°C, 5min at a speed of 370g for most washing procedures.

### Naïve CD4^+^ T cell isolation

Naïve CD4^+^ T cells were isolated from cell suspensions with pooled lymph nodes and spleen. Isolation was performed with StemCell mouse naïve CD4^+^ T cell isolation kit according to manufacturer’s instructions. In brief, the cell suspensions were incubated with rat serum and CD4^+^ isolation antibody for 7.5 minutes, then with memory depletion antibody for 2.5 minutes, and in the end with magnetic beads for another 2.5 minutes before incubating with the isolation magnet for 2.5 minutes. The resulting untouched naïve CD4^+^ T cells were then washed with FACS buffer, and purity was routinely checked with a staining for CD4^+^, CD44^-^ and viability.

### Plate-bound CD4^+^ T cell activation

Plates were coated over night with 0.2μg anti-CD28 and 0.5μg anti-CD3 per ml PBS for most of the experiments (low stimulation as according to [1]). We used 1ml/well for 24 well plates and 0.2ml/well for 96 well plates. Before plating of the cells, plates were washed with PBS. We plated 2*10^5^ naïve CD4^+^ T cells per well in 96 well flat-bottom in 200μl medium. For 24 well plates, 2*10^6^ naïve CD4^+^ T cells per ml medium in 1ml medium were plated. Complete T cell medium (RPMI-1640 Medium, 10% FCS, 1% HEPES, 1% non-essential amino acids, 1% Glutamax, 0.1% 2-Mercaptoethanol) was supplied with 50U IL-2/ml. Cells were cultured at 37°C, 5% CO_2_ for 24h or longer depending on the purpose of the experiment as indicated in figure legends.

### Proliferation assay with cell trace violet (CTV)

Freshly isolated naïve CD4^+^ T cells were washed with PBS. 1μl of Cell Trace stock solution (dissolved in DMSO according to the manufacturer’s instructions) was then used per ml PBS for 10*10^6^ cells. Cells were incubated at 37°C for 20 minutes, then 5x the original staining volume of normal T cell culture medium was added for 5 minutes to remove residual dye. Cells were washed and plated in complete culture medium supplied with 50U IL-2 per ml for 48h.

### Enzyme-linked immunosorbent assay (ELISA)

IL-2 secretion was addressed in 48h culture supernatants from cells that were plated in precoated wells in complete T cell medium without IL-2. IL-2 ELISA was performed with the BioLegend ELISA MAX mouse IL-2 set according to the manufacturer’s instructions. After the last washing step, TMB substrate was added for the readout. Absorbance was measured with an ELISA plate reader (Synergy H1 Hybrid Reader, BioTek) at 450nm as well as 570nm wavelength, and normalized to wild type control samples that were run in the same experiment.

### FACS Staining

Generally, cells were stained for viability with viability dye 780 in PBS for 20min at 4°C and then washed with PBS. Non-specific binding was blocked with anti-CD16/anti-CD32 0.5mg/ml on ice for 10 minutes. Surface staining was performed in FACS buffer for 20-30 minutes at 4°C. In intracellular staining or LCMV experiments, cell fixation was performed with Fix-Perm for 20 minutes on 4°C (1h for LCMV experiments). Intracellular staining was done in permeabilization buffer for 1h at 4°C. Activation status of the cells was assessed by staining and gating for singlets/lymphocytes/viable cells/ CD4^+^ and early activation marker CD25/CD69 as well as CD44/CD62L expression. Blasting of lymphocytes was addressed by pre-gating on singlets/viable cells. For cytokine staining after *in vitro* differentiation or *ex vivo* e.g., for IL-2 staining, cells were stimulated with 50ng/ml PMA, 500ng/ml Ionomycin and 10 μg/ml Brefeldin A (BFA) for 3h at 37°C before staining. Data was acquired with an LSR Fortessa (BD) and analyzed with FlowJo (version 10.4.1).

### LCMV Armstrong infection model

Mice were infected with 2*10^5^ PFU LCMV-Armstrong strain i.p. with U-100 insulin syringes (0.30mm (30G) x 8mm). Eight days post infection, the animals were euthanized with CO_2_ and the spleens were harvested for staining. We gated for singlets/lymphocytes/viable cells/CD3^+^CD4^+^ to look at CD44 expression, and moreover we addressed T_FH_ cells by the coexpression of key markers Bcl-6, PD-1, ICOS and CXCR5. We gated for singlets/lymphocytes/viable cells/CD19^+^B220^+^ to look at GC B cells expressing Fas and GL-7. Re-stimulation of splenocytes was performed in flat bottom 96 well plates with 1μg/ml LCMV-specific peptide GP-64 in comparison to polyclonal 50ng/ml PMA, 500ng/ml Ionomycin stimulation for one hour, then 10μg/ml Brefeldin A was added for another three hours before staining. We then gated for T_H_1 cells using singlets/lymphocytes/viable cells/CD3^+^CD4^+^ and finally looking at Tbx21/IFNγ expression. All in vivo experiments were performed blinded with assignment of animal numbers after data analysis.

### Histology

Spleens were embedded in cryo embedding medium and frozen on dry ice before storage at −80°C. Sections were cut at a thickness of 6μm and dried on air. Single sections were then fixed with acetone for 5 minutes and circled with PAP pen. Staining for CD19, CD4 and GL-7 was performed in FACS buffer with anti-CD16/anti-CD32 in a wet chamber at 4°C overnight. Slides were then washed with PBS on a shaker for 15 minutes before drying and mounting. Imaging was performed with a 20x objective on a Nikon Ti2 microscope and analyzed with ImageJ version 2.0.0.

### Adoptive transfer (AT) with subsequent LCMV infection

Naïve SMARTA^+^ CD4^+^ T cells from wt, CD28^-/-^ and rescue mice were isolated transferred into CD28^-/-^ recipients. Each recipient received 1*10^5^ cells in 100μl PBS i.v.. The recipients were infected with 2*10^5^ PFU LCMV Armstrong i.p. two days after cell transfer. Eight days after infection, the mesenteric and peripheral LN as well as the spleen were analyzed separately for the presence of Vα2^+^Vβ8.3^+^ T cells (pre-gated on singlets/lymphocytes/viable cells/CD4^+^), and these cells were further characterized for their CD44 expression. One CD28^-/-^ mouse that did not receive donor cells was used as a negative control in each experiment to display the recipient-intrinsic Vα2^+^Vβ8.3^+^ population.

### RNA sequencing

For any experiment involving RNA, cells were washed with PBS before counting and the RNA was kept on ice during the experiments, storage at −80°C. All pipetting was performed with filter-tips and RNAse-free tubes. All procedures for the extractions were performed at the facilities with materials, protocols and supervision of the facility experts. For RNA sequencing, 2.5*10^5^ cells were washed with PBS and resuspend in 200μl TRI Reagent. RNA was extracted from Trizol-samples with a Zymo Direct-zol kit which includes DNAse treatment. Quality control was run with a Bioanalyzer. RNA quality was assessed with a Fragment Analyzer (Advanced Analytical) and RNA-seq library preparation was performed using Illumina Truseq stranded kit. Sequencing was performed on an Illumina NexSeq 500 machine to produce single-end 76-mers reads. All steps were performed at the Genomics Facility Basel (ETH Zurich).

Read quality was assessed with the FastQC tool (version 0.11.5). Reads were mapped to the mouse genome (UCSC version mm10) with STAR (version 2.5.2a) [2] with default parameters, except filtering out reads mapping to more than 10 genomic locations (outFilterMultimapNmax=10), reporting only one hit in the final alignment for multimappers (outSAMmultNmax=1) and filtering reads without evidence in the spliced junction table (outFilterType-“BySJout”).

All subsequent gene expression data analysis was performed using the R software (version 3.5). Read alignment quality was evaluated using the qQCReport function of the R Bioconductor package QuasR (version 1.18). Gene expression was quantified using the qCount function of QuasR [3] as the number of reads (5’ends) overlapping with the exons of each gene assuming an exon union model (using the UCSC knownGenes annotation downloaded on 2015-12-18). To quantify intronic expression levels, exonic coordinates were extended by 10 bp on each side of the exons, and for each gene the resulting read count was subtracted to the read count obtained on the whole gene (extended by 10 bp on each side).

The R Bioconductor package edgeR (version 3.28)[4] was used for differential gene expression analysis. Between samples normalization was done using the TMM method [5]. Only genes with CPM (counts per million reads mapped) values more than 1 in at least 4 samples (the number of biological replicates) were retained. Principal component analysis (PCA) was performed on the log-transformed normalized CPM values. An generalized linear model including a genotype effect, an activation effect, and a replicate effect (nested within genotype) was fitted to the raw counts (function glmFit), and differential expression was tested using likelihood ratio tests (function glmLRT). P-values were adjusted by controlling the false-discovery rate (Benjamini-Hochberg method) and genes with a FDR lower than 1% were considered differentially expressed.

To select a restricted list of most likely miRNA targets for each seed family, we restricted the genes to those with a conserved 3’UTR seed match (using TargetScan [6]), and a coverage of more than 5 AHC reads. We additionally restricted the list by using differential expression criteria from the first RNA-seq dataset: genes should be significantly differentially expressed between T^1792tg/tg^ and wt, as well as between T^1792Δ/Δ^ and wt, with fold-changes in the expected directions; using intronic read counts, genes should not be significantly differentially expressed in the same comparison, at an increased FDR threshold of 10%.

Gene set enrichment analysis was performed with the function camera [7] from the limma package (version 3.42; using the default parameter value of 0.01 for the correlations of genes within gene sets) using gene sets from the curated gene set collection (C2) of the Molecular Signature Database (MSigDB v7.0)[8] with a special focus on gene sets from pathway databases, including KEGG [9], Biocarta (http://cgap.nci.nih.gov/Pathways/BioCarta_Pathways), PID [10] and Reactome [11]. We tested a total of 1,576 gene sets containing more than 10 genes and those with a false discovery rate (FDR) lower than 1% were considered significant.

### Regulon enrichment analysis

Regulon enrichment analysis was performed similarly to pathway enrichment analysis using camera from the limma package. DoRothEA v2 regulons [11] were downloaded from https://github.com/saezlab/DoRothEA.

More specifically, we used human TOP10score regulons in VIPER format, containing the targets with the highest quality score possible (and at least 10 genes) per transcription factor. Mouse regulons were obtained by orthology: mouse genes with a 1-to-1 or Many-to-1 relationship to human genes of each DoRothEA regulon were obtained using Ensembl Compara (release 97); only orthology relationships annotated to taxonomic levels “Boreoeutheria”, “Eutheria”, “Euarchontoglires”, “Theria”, or “Amniota” were retained. We tested a total of 1,307 regulons including at least 10 mouse genes and those with a false discovery rate (FDR) lower than 1% were considered significant.

### AGO2 HITS-CLIP (AHC) data

Read depths on seed regions of genes targeted by different miRNA seed families were obtained as previously described [12].

### In vitro differentiation

T_H_1 differentiation conditions were generated with 50U IL-2, 5ng/ml IL-12 and 10μg/ml anti-IL-4 per ml T cell medium [13]. iTregs were either differentiated with retinoic acid (0.9mM), 250U IL-2, 0.75ng/ml TGFβ, 10μg/ml anti-IFNγ and 10μg/ml anti-IL-4 [14]. T_H_17 were generated with 50ng/ml IL-6, 3ng/ml TGFβ, 5μg/ml anti-IFNγ and 10μg/ml anti-IL-4 per ml T cell medium [15]. For the differentiation, 2*10^5^ naïve CD4^+^ T cells were plated on a precoated 96-well flat bottom plate (coated over night with 0.2μg anti-CD28, 0.5μg anti-CD3 per ml PBS) and harvested at 24h, 48h or 72h after plating.

### Seahorse

Calibration plates were coated over night with 200μl calibrant. Cell plates were coated with 18μl 0.1M NaHCO_3_ pH8.0 6.67% CellTak (Seahorse XF96 flux pack, Bucher Biotech, CH). The following day, cell plates were washed with H_2_O and left for drying during cell preparation. Compounds were prepared for a final in-well concentration of 1μM for Oligomycin, 2μM for FCCP and 11μM for Rotenone. CD4^+^ T cells (naïve or activated) were harvested with glucose-free, unbuffered RPMI, washed and counted multiple times before plating 3*10^5^ cells per well. Mitochondrial perturbation was performed by sequential injection of glucose (80mM stock), oligomycin, FCCP and rotenone. Measurements of oxygen consumption rate (OCR, pMoles/min) and extracellular acidification rate (ECAR; mpH/min) were performed with a Seahorse XF96 flux analyzer (Seahorse Bioscience, USA). Data analysis was performed using Prism (Version7.0d), mitochondrial parameters were calculated as described by Gubser et al. [16].

### CyA titration

The sensitivity of CD28^-/-^ and rescue cells to compounds interfering with Ca^2+^ signaling was tested. Increasing amounts of CyA were added to the cultures during activation: 2*10^5^ naïve CD4^+^ T cells were plated in 100μl complete T cell medium in pre-coated 96 well plates. 100μl of a serial dilution of CyA were then added to result in in-well concentrations of 100ng/ml, 50ng/ml, 25ng/ml, 12.5ng/ml, 6.5ng/ml, 3.125ng/ml, 1.5625ng/ml or 0ng/ml. Cells were then activated in the presence of these CyA concentrations for 48h before harvesting and staining for viability and activation markers.

### ImageStream

Naïve CD4^+^ T cells were isolated and activated for 48h in the presence of 6.25ng/ml CyA. They were then harvested and washed with PBS before fixation for 20min at 4°C. Intracellular staining for NFATc2 was performed in a two-step staining with first 1h at room temperature with anti-NFATc2 in permeabilization buffer, and subsequently 1h at room temperature with goat anti-mouse IgG1 in permeabilization buffer. Nuclei were stained with DAPI in the last 5min of incubation. Acquisition was run on an ImageStreamX Mark2 Imaging flow cytometer (Amnis), and data analysis was performed with the IDEAS software (v6.2).

### Reverse transcription (RT) and quantitative PCR (qPCR)

5*10^5^ cells were washed resuspend in 400μl TRIreagent. RNA isolation was then performed according to the isolation protocol from TRIreagent supplier (SIGMA). In brief, 0.1ml of 1-bromo-3-chloropropane per ml of TRI Reagent was added, samples were mixed by vigorous shaking, incubated for 15min at room temperature and then centrifuged at 12’000g for 15min at 4°C for phase separation. The aqueous phase was then mixed with 0.5 ml of isopropanol per ml of TRI Reagent used, again centrifuged for 10min for RNA precipitation. RNA was then washed with 70% Ethanol and finally resuspend in RNAse-free water. RNA concentration and purity was determined with a Nanodrop2000 Spectrophotometer (ThermoScientific).

Reverse transcription was performed with RNA extracted with TRIzol and the SIGMA MMLV kit on 1μg RNA according to the manufacturer’s instructions. qPCR was run with TaqMan FAST Universal PCR master mix on an Applied Biosystems® Real-Time PCR System. 18S was used as a reference gene.

### Proteomics

All procedures for the extractions were performed at the facilities with materials, protocols and supervision of the facility experts. Heavy labeled reference peptides for Rcan3 identification were ordered from JPT (JPT.com, Berlin. Germany). For targeted proteomics, 2.5*10^5^ cells were washed with PBS and pellets were frozen on dry ice until protein extraction. Cells were then lysed in lysis buffer (1% Sodium deoxycholate, 10mM TCEP, 100mM Tris, pH=8.5 adjusted with NaOH/HCl) for 1 minute at 50°C, shaking at 300rpm. Samples were then sonicated (30 sec on, 30 sec off, 10 cycles, Bioruptor, Diagnode) and heated for 10 minutes, 95°C at 300rpm. Chloroacetamide (0.75M) was added to a final concentration of 15mM and the protein samples incubated for 30 minutes at 37°C, shaking at 500rpm. pH was then adjusted to 8-9 using 1M Ammoniumbicarbonate where necessary. Proteins were then digested using Lysin-C (enzyme/protein ratio of 1:200, 37°C for 3h) and trypsin (enzyme/protein ratio of 1:50, 37°C overnight). Peptide samples were then centrifuged at 5000rpm before adding 5% TFA and washing in wash buffer (1% TFA in 2-propanol). Peptides were cleaned up using the Phoenix 96× kit following the vendor protocol (PreOmics, Martinsried, Germany). For LC-MS/MS analysis, peptides were dissolved in LC buffer A (0.15% Formic acid, 2% acetonitrile, shake at 1,400 rpm at 25°C for 5 minutes) to a final peptide concentration of 0.5 μg/μl. iRT-peptides mix was added before analysis.

For LC-MS analysis, parallel reaction-monitoring (PRM) assays [17] were generated from a mixture containing 100 fmol of each heavy reference peptide and shotgun data-dependent acquisition (DDA) LC-MS/MS analysis on a Thermo Orbitrap Fusion Lumos platform (Thermo Fisher Scientific). The setup of the μRPLC-MS system was as described previously [18]. Chromatographic separation of peptides was carried out using an EASY nano-LC 1000 system (Thermo Fisher Scientific), equipped with a heated RP-HPLC column (75 μm x 30 cm) packed in-house with 1.9 μm C18 resin (Reprosil-AQ Pur, Dr. Maisch). Peptides were analyzed per LC-MS/MS run using a linear gradient ranging from 95% solvent A (0.15% formic acid, 2% acetonitrile) and 5% solvent B (98% acetonitrile, 2% water, 0.15% formic acid) to 45% solvent B over 60 minutes at a flow rate of 200 nl/min. Mass spectrometry analysis was performed on Thermo Orbitrap Fusion Lumos mass spectrometer equipped with a nanoelectrospray ion source (both Thermo Fisher Scientific). Each MS1 scan was followed by high-collision-dissociation (HCD) of the 10 most abundant precursor ions with dynamic exclusion for 20 seconds. Total cycle time was approximately 1 s. For MS1, 1e6 ions were accumulated in the Orbitrap cell over a maximum time of 100 ms and scanned at a resolution of 120,000 FWHM (at 200 m/z). MS2 scans were acquired at a target setting of 1e5 ions, accumulation time of 50 ms and a resolution of 30,000 FWHM (at 200 m/z). Singly charged ions and ions with unassigned charge state were excluded from triggering MS2 events. The normalized collision energy was set to 30%, the mass isolation window was set to 1.4 m/z and one microscan was acquired for each spectrum.

The acquired raw-files were database searched against a *Mus musculus* database (Uniprot, download date: 2017/04/18, total of 34,490 entries) by the MaxQuant software (Version 1.0.13.13). The search criteria were set as following: full tryptic specificity was required (cleavage after lysine or arginine residues); 3 missed cleavages were allowed; carbamidomethylation (C) was set as fixed modification; Arg10 (R), Lys8 (K) and oxidation (M) as variable modification. The mass tolerance was set to 10 ppm for precursor ions and 0.02 Da for fragment ions. The best 6 transitions for each peptide were selected automatically using an in-house software tool and imported to SpectroDive (v8.0, Biognosys, Schlieren, Switzerland). A mass isolation lists containing all selected peptide ion masses were exported and imported into the QE-HF operating software for PRM analysis using the same LC and MS setting as above with the following modifications: The resolution of the orbitrap was set to 120k FWHM (at 200 m/z) and the fill time was set to 250 ms to reach a target value of 3e6 ions. Ion isolation window was set to 0.4 Th and the scan range was set to 100-1500 Th. Normalized collision energy was set to 27%. A MS1 scan using the same conditions are for DDA was included in each MS cycle. All raw-files were imported into SpectroDive for protein / peptide quantification. To control for variation in sample amounts, the total ion chromatogram (only comprising peptide ions with two or more charges) of each sample was determined by Progenesis QI (version 2.0, Waters) and used for normalization.

### Quantification and statistical analysis

Statistical analysis was performed with GraphPad Prism (v7.0d, Graphpad Software). Normal distribution was not assumed, and non-parametric tests were chosen individually depending on the type of experiment as indicated in the figure legends. The overall statistical significance was set to 5% (α=0.05), and error bars represent standard deviation (SD). N corresponds to number of biological replicates. We used Kruskal-Wallis tests for most of the experiments with more than two unpaired groups and one factor (e.g., %Fas^+^GL7^+^ population in three genotypes), followed by Dunn’s multiple comparison test. We used two-way Anova for experiments including two factors (e.g., %CD44^hi^CD62^lo^ population in three genotypes activated with or without anti-CD28), followed by Tukey’s multiple comparison test. A *t*-test was used for the analysis of targeted proteomics (each genotype compared to wt).

