## Supplemental Figures and Tables for "The non-coding RNA miR-17~92 is a central mediator of T cell activation"

**Supplementary Material:**

**Supplementary Figures S1-S7**
**Supplementary Tables 1 and 2**

### The non-coding RNA miR-17~92 is a central mediator of T cell activation

**Authors**

Marianne Dölz[1, 2], John D. Gagnon[3, 4, 5], Mara Kornete[1, 6], Romina Marone[1, 2], Glenn Bantug[1], Robin Kageyama[3, 4], Christoph Hess[1, 7], K. Mark Ansel[3,4], Denis Seyres[1, 2], Julien Roux[1,8] and Lukas T. Jeker[1, 2]*

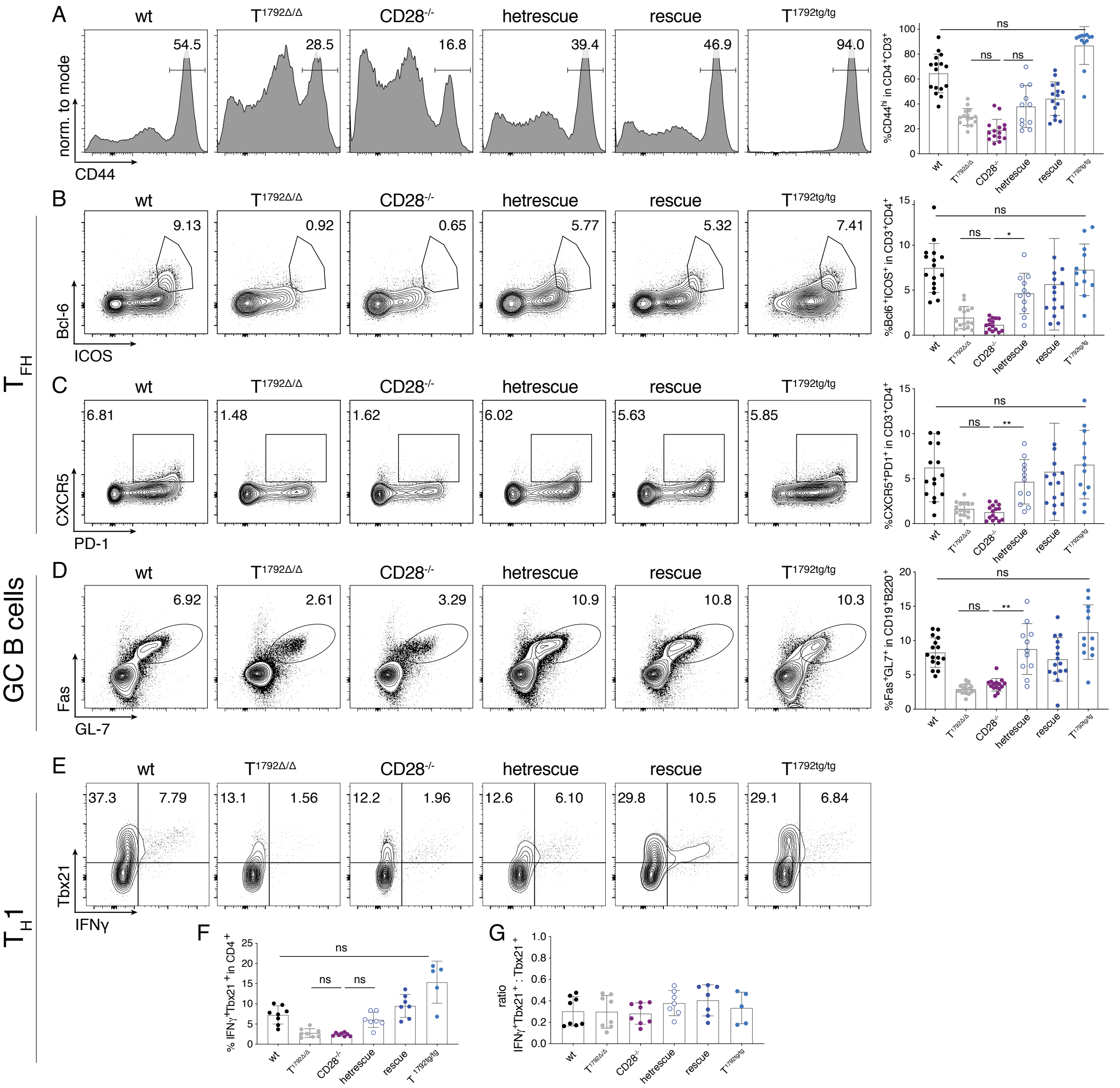


**Supplemental Figure 1 rel. to Figure 3. miR-17~92-mediated gene regulation is very sensitive to precise miR-17~92 expression in vivo.**

6-8 week old mice were infected with 2*105 PFU LCMV Armstrong and spleens were analyzed at d8 post infection. wt (black), T1792Δ/Δ (grey), CD28-/- (purple), CD28-/- with heterozygous transgenic miR-17~92 expression (hetrescue; dark blue empty circles), CD28-/- with homozygous transgenic miR-17~92 expression (rescue; dark blue), T1792tg/tg (light blue). A-D) represent data from 4 independent experiments with 3-4 mice per group, gated on viable CD4+CD3+ or viable CD19+B220+ cells. A) CD44 expression B) quantification of % Bcl6+ICOS+ (TFH) C) % CXCR5+PD-1+ (TFH) and D) % Fas+GL7+ (GC B cells). Error bars represent mean with SD, Dunn's multiple comparison test, p values: ns=not significant, *<0.0332, **<0.0021, ***<0.0002, ****<0.0001. E-G) Splenocytes were restimulated with GP-64 and BFA for 4h and investigated for TH1 phenotype, pre-gated on viable CD3+CD4+ cells. Shown are 2-3 independent experiments with 3-4 biological replicates per group. E) representative contour plots of Tbx21 and IFNγ expression F) Tbx21+IFNγ+ data summary G) ratio of Tbx21+IFNγ+ to total Tbx21+ cells.

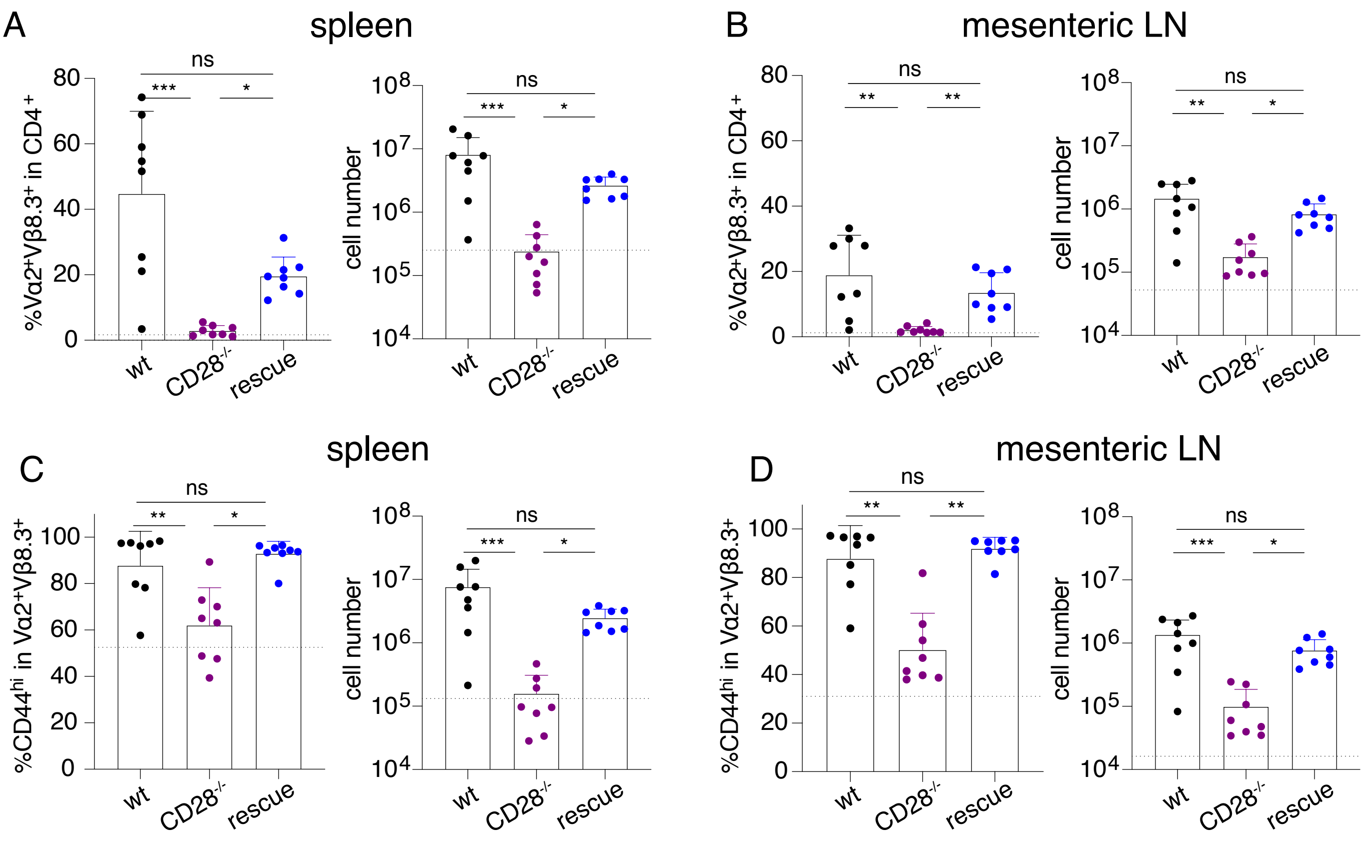


**Supplemental Figure 2, rel. to Figure 4. Restoration of T cell activation of CD28-deficient T cells by miR-17~92 is cell intrinsic.**

AT of naïve SMARTA+ CD4+ T cells into CD28-/- hosts, subsequent LCMV Armstrong infection and analysis of organs at d8 post infection. Donor genotypes wt (black), CD28-/- (purple), rescue (dark blue). Dotted line indicates recipient's intrinsic Vα2+Vβ8.3+ population measured in a non-transferred host. Vα2+Vβ8.3+ cells in viable CD4+ population from spleen (A) and mesenteric LN (B). CD44 expression in Vα2+Vβ8.3+ population from spleen (C) and mesenteric LN (D). 2 independent experiments, 4 recipients per group. Error bars represent mean ±SD, Dunn's multiple comparison test, p values: ns=not significant, *<0.05, **<0.002, ***<0.0002.

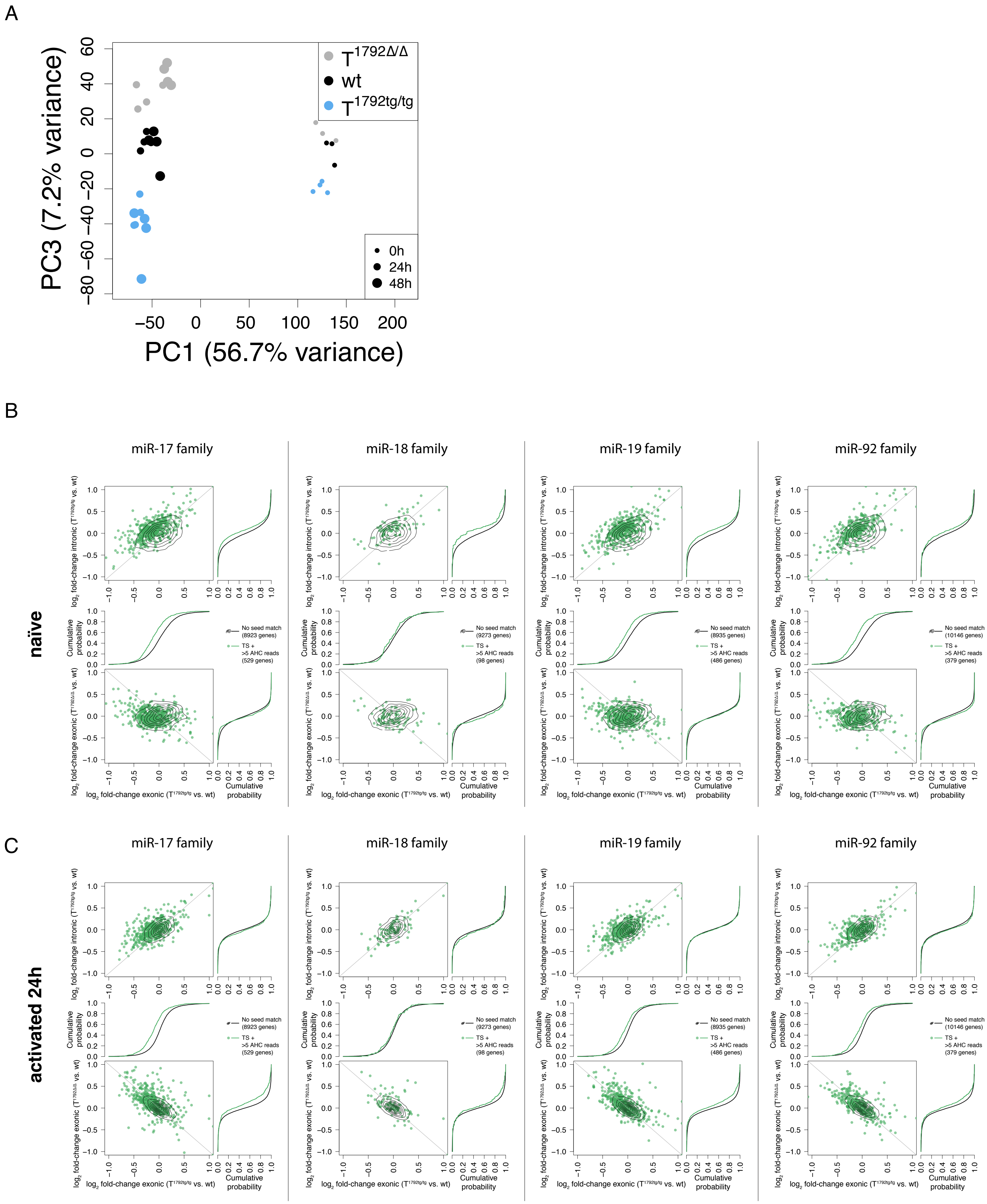


**Supplemental Figure 3 rel. to Figure 5. miR-17~92 shapes the transcriptome after CD4+ T cell activation**

Naïve CD4+ T cells from T1792Δ/Δ (grey), wt (black) and T1792tg/tg (light blue) mice were activated with plate-bound αCD28 and αCD3 mAbs for 0, 24h and 48h. Total RNA was extracted for bulk sequencing. A) PCA (PC1 vs. PC3) based on the 25% most variable genes.B, C) For each seed family, dot plots compare the log2 value of the exonic expression ratio for each gene in T1792tg/tg vs. wt (x-axis) versus either exonic expression ratio for each gene in T1792Δ/Δ vs. wt comparison on the y-axis (bottom row) or intronic expression ratio for each gene in T1792tg/tg vs. wt (top row) in naïve (B) and 24h post activation (C). Each ratio is compared to the cumulative fraction of all log2 ratios in the corresponding comparison. Black curve: all genes of our data set without a seed match and ≤5 AHC reads; green: subset of genes with a seed sequence for the indicated seed family and >5 reads in the AHC.

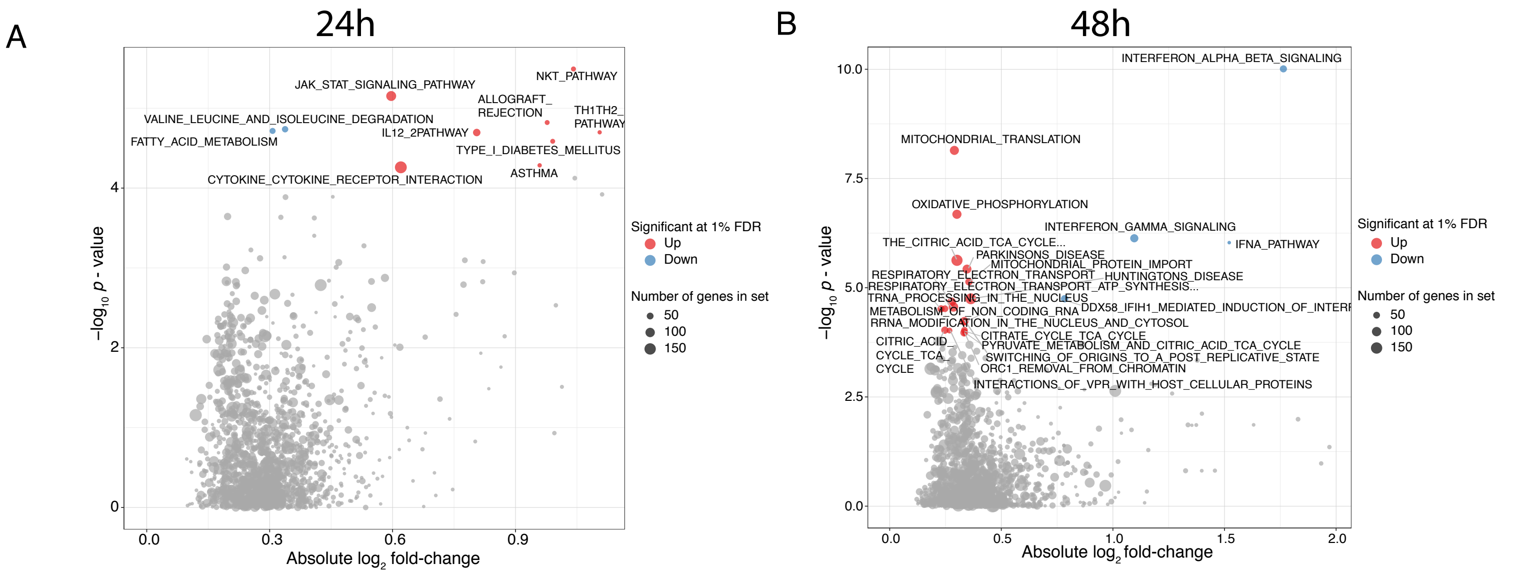


**Supplemental Figure 4 rel. to Figure 6. GSEA pathway analysis at 24h and 48h**
Naïve CD4+ T cells from T1792Δ/Δ , wt and T1792tg/tg mice were activated with plate-bound αCD28 and αCD3 mAbs for 0 and 24h. Total RNA was extracted for sequencing. Volcano plot of GSEA pathway (KEGG,REACTOME,PID,BIOCARTA) enrichment for genes differentially expressed after 24h (A) and 48h (B) activation between T1792Δ/Δ  and T1792tg/tg. Red dots indicate pathway enriched for genes up regulated whereas blue dots are pathways enriched for down regulated genes. Only significantly enriched pathways (1% FDR) are colored. Dot width indicates the gene set size for each pathway.

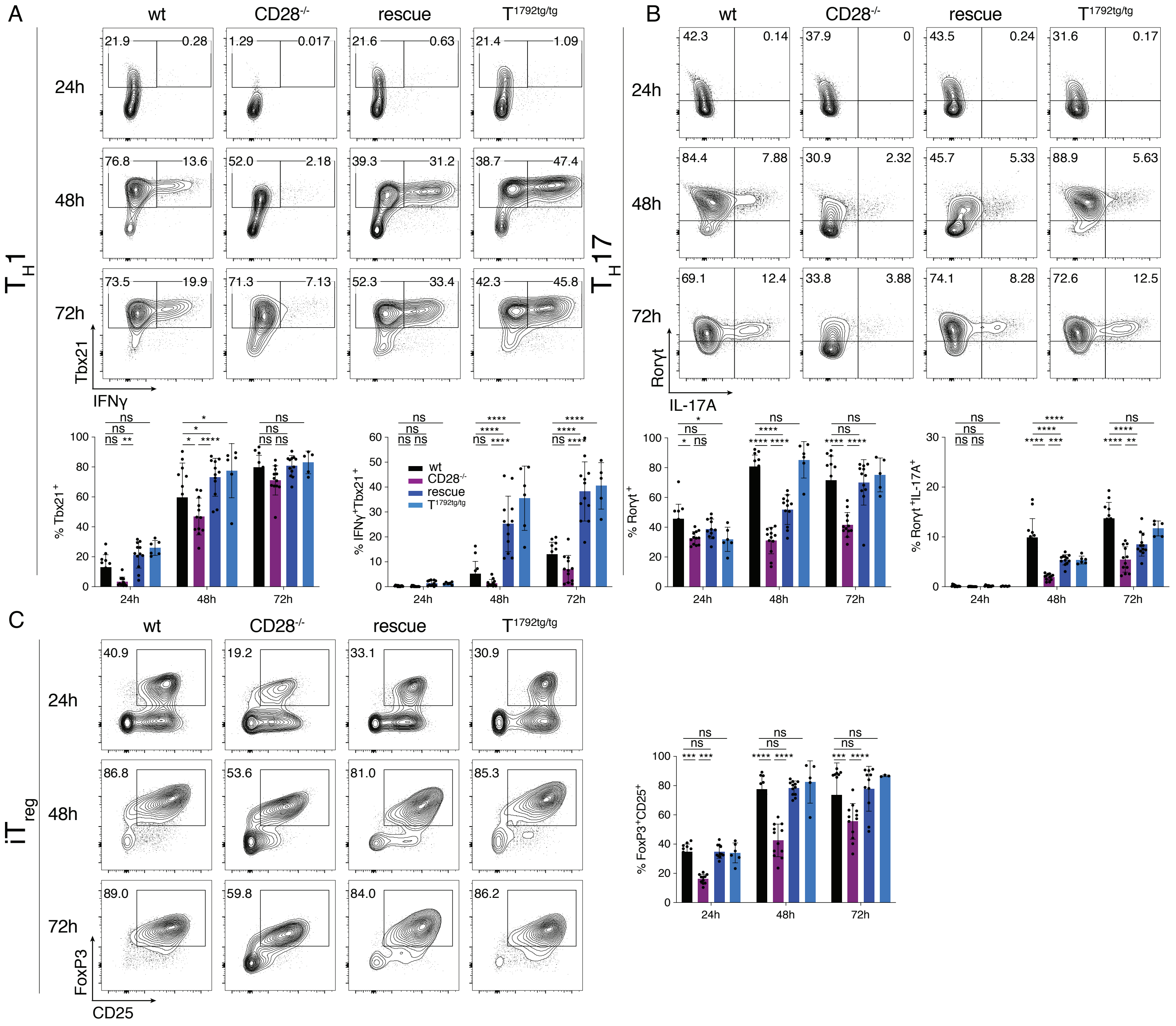

**Supplemental Figure 5 rel. to Figure 6. Transgenic miR-17~92 functionally corrected the defects of CD28-/- T cells to differentiate into TH1, TH17 and iTreg in vitro.**

Naïve CD4+ T cells were activated for 24h, 48h, and 72h with plate-bound αCD28 and αCD3 under in vitro skewing conditions for TH1, TH17, and iTreg differentiation. wt (black), CD28-/- (purple), rescue (dark blue) and T1792tg/tg (light blue). A) TH1 differentiation assessed by IFNγ/Tbx21 staining in viable CD4+ T cells B) TH17 differentiation assessed by IL-17A/Rorγt staining in viable CD4+ T cells C) iTregs characterized by CD25/ Foxp3 expression. Data from 2 independent experiments.

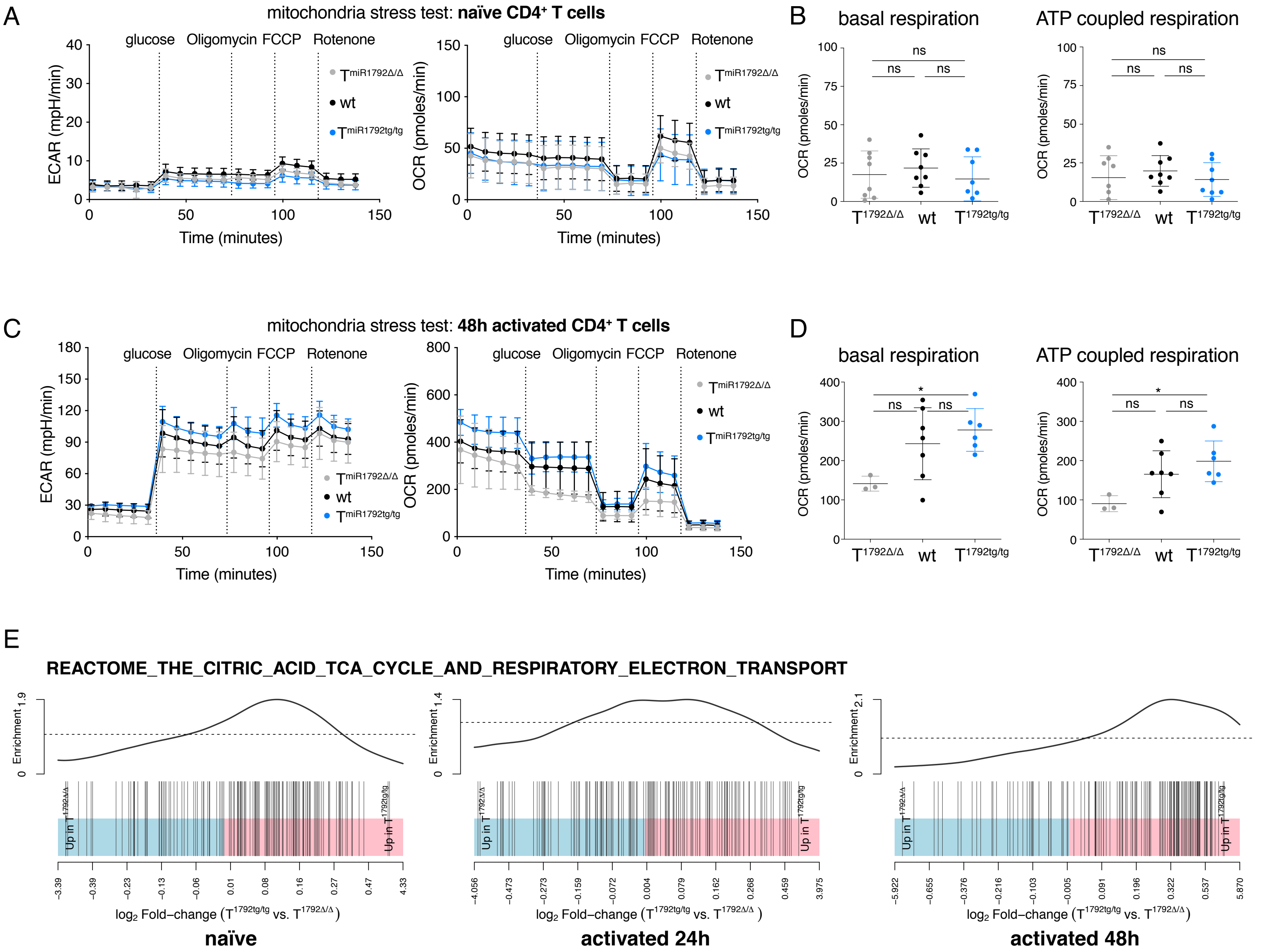


**Supplemental Figure 6 rel. to Figure 6. miR-17~92 expression modifies metabolism in activated CD4+ T cells after 48h**
CD4+ T cells from T1792Δ/Δ (grey), wt (black), T1792tg/tg (light blue) were assessed for their metabolic activity by mitochondria stress test with the seahorse machine. A) Mitochondria stress test measured with a 96-well seahorse in naïve CD4+ T cells. 2 experiments with 3-4 biological replicates per group and experiment are shown. B) Basal respiration and ATP coupled respiration in naïve CD4+ T cells C) Mitochondria stress test measured in CD4+ T cells activated for 48h. D) Basal respiration and ATP coupled respiration in activated CD4+ T cells. Pooled 3-8 biological replicates from 4 experiments are shown. Tukey's multiple comparison test, p values: *<0.05. E) RNA sequencing data shows an enrichment of genes associated with TCA. Shown is gene set REACTOME_TCA_CYCLE_AND_RESPIRATORY_ ELECTRON_TRANSPORT enrichment with genes differentially expressed between T1792Δ/Δ and T1792tg/tg in naïve, 24h and 48h post activation. Colors indicate fold change direction with blue being upregulated in T1792Δ/Δ and red downregulated.

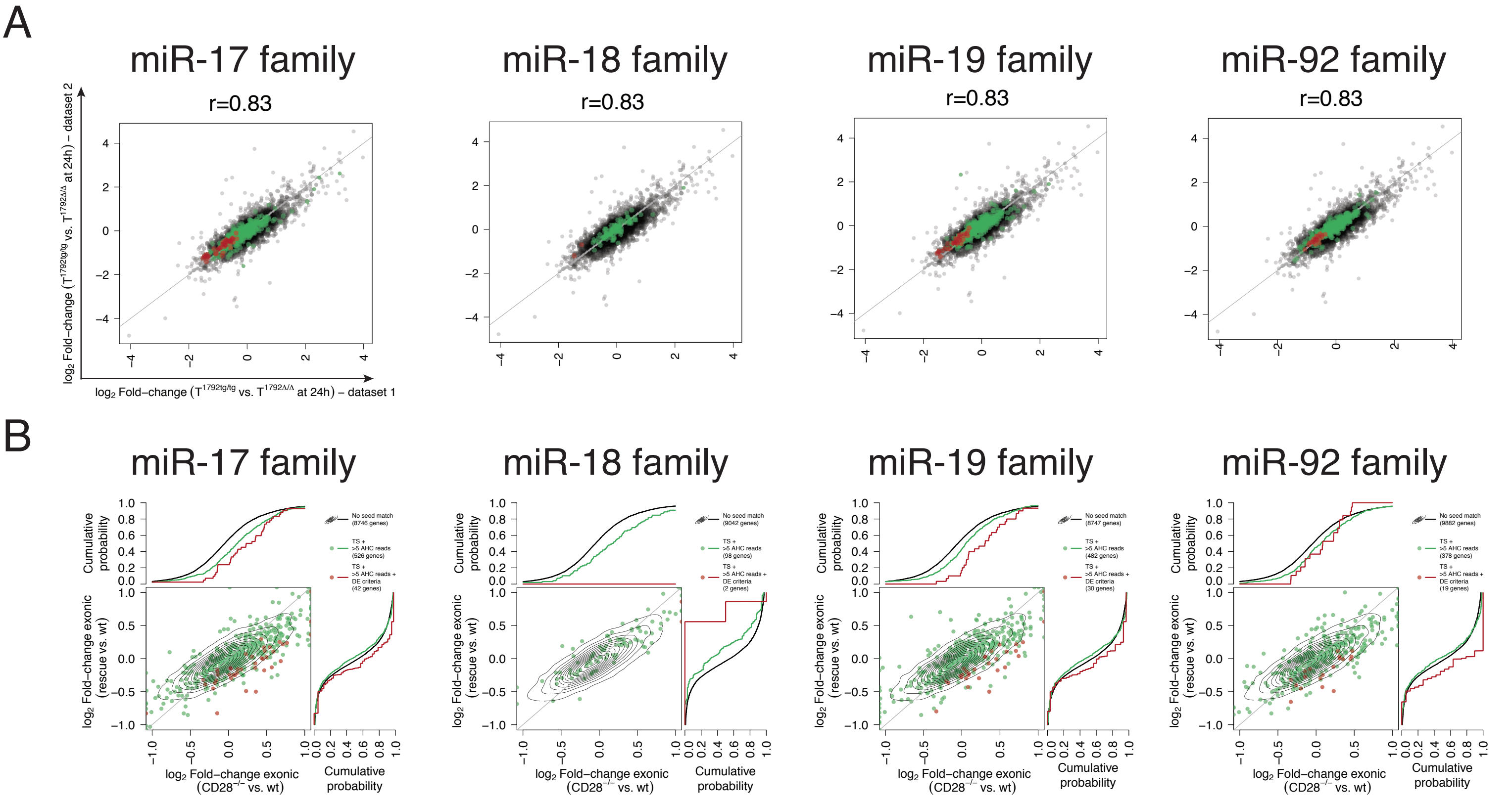


**Supplemental Figure 7 rel. to Figure 7. All miR-17~92 seed families are necessary for CD28-induced gene repression**
Total RNA was extracted for sequencing from CD4+ T cells. A) Correlation of gene expression levels between RNA-seq experiment 1 (Fig. 5) and 2 (Fig. 7). Black: genes without seed match, <5 AHC reads and no differential expression in the second RNA sequencing. Green: genes with a conserved binding site for each indicated seed family and >5 reads in the AHC; red: genes with a conserved binding site for each indicated seed family, >5 reads in the AHC and differential expression in the second RNA sequencing data set. B) For each seed family, dot plots compare the log2 value of the exonic expression ratio for each gene in the CD28-/- vs. wt comparison (x-axis) versus the exonic expression ratio for each gene in rescue vs. wt (y-axis) in CD4+ T cells 24h post activation. Each ratio is compared to the cumulative fraction of all log2 ratios in the corresponding comparison. Black curve: all genes of our data set without a seed match and five or less AHC reads; green: subset of genes with a seed sequence for the indicated seed family (TS) and >5 reads in the AHC; red: genes with a conserved binding site for the corresponding seed family (TS), >5 reads in the AHC and differential expression in the second RNA sequencing data set.

**Supplementary Figure legends**

**Supplemental Figure 1 rel. to Figure 3**
**miR-17~92-mediated gene regulation is very sensitive to precise miR-17~92 expression in vivo**
6-8 week old mice were infected with $2*10^5$ PFU LCMV Armstrong and spleens were analyzed at d8 post infection. wt (black), $T^{1792\Delta/\Delta}$ (grey), $CD28^{-/-}$ (purple), $CD28^{-/-}$ with heterozygous transgenic miR-17~92 expression (hetrescue; dark blue empty circles), $CD28^{-/-}$ with homozygous transgenic miR-17~92 expression (rescue; dark blue), $T^{1792tg/tg}$ (light blue). A-D) represent data from 4 independent experiments with 3-4 mice per group, gated on viable $CD4^+CD3^+$ or viable $CD19^+B220^+$ cells. A) CD44 expression B) quantification of % $Bcl6^+ICOS^+$ ($T_{FH}$) C) % $CXCR5^+PD\text{-}1^+$ ($T_{FH}$) and D) % $Fas^+GL7^+$ (GC B cells). Error bars represent mean with SD, Dunn's multiple comparison test, p values: ns=not significant, *<0.0332, **<0.0021, ***<0.0002, ****<0.0001. E-G) Splenocytes were restimulated with GP-64 and BFA for 4h and investigated for $T_H1$ phenotype, pre-gated on viable $CD3^+CD4^+$ cells. Shown are 2-3 independent experiments with 3-4 biological replicates per group. E) representative contour plots of Tbx21 and IFNγ expression F) $Tbx21^+IFN\gamma^+$ data summary G) ratio of $Tbx21^+IFN\gamma^+$ to total $Tbx21^+$ cells.

**Supplemental Figure 2, rel. to Figure 4**
**Restoration of T cell activation of CD28-deficient T cells by miR-17~92 is cell intrinsic**

AT of naïve $SMARTA^+$ $CD4^+$ T cells into $CD28^{-/-}$ hosts, subsequent LCMV Armstrong infection and analysis of organs at d8 post infection. Donor genotypes wt (black), $CD28^{-/-}$ (purple), rescue (dark blue). Dotted line indicates recipient's intrinsic $V\alpha2^+V\beta8.3^+$ population measured in a non-transferred host. $V\alpha2^+V\beta8.3^+$ cells in viable $CD4^+$ population from spleen (A) and mesenteric LN (B). CD44 expression in $V\alpha2^+V\beta8.3^+$ population from spleen (C) and mesenteric LN (D). 2 independent experiments, 4 recipients per group. Error bars represent mean ±SD, Dunn's multiple comparison test, p values: ns=not significant, *<0.05, **<0.002, ***<0.0002.

**Supplemental Figure 3 rel. to Figure 5**
**miR-17~92 shapes the transcriptome after $CD4^+$ T cell activation**

Naïve $CD4^+$ T cells from $T^{1792\Delta/\Delta}$ (grey), wt (black) and $T^{1792tg/tg}$ (light blue) mice were activated with plate-bound αCD28 and αCD3 mAbs for 0, 24h and 48h. Total RNA was extracted for bulk sequencing. A) PCA (PC1 vs. PC3) based on the 25% most variable genes.
B, C) For each seed family, dot plots compare the log2 value of the exonic expression ratio for each gene in $T^{1792tg/tg}$ vs. wt (x-axis) versus either exonic expression ratio for each gene in $T^{1792\Delta/\Delta}$ vs. wt comparison on the y-axis (bottom row) or intronic expression ratio for each gene in $T^{1792tg/tg}$ vs. wt (top row) in naïve (B) and 24h post activation (C). Each ratio is compared to the cumulative fraction of all log2 ratios in the corresponding comparison. Black curve: all genes of our data set without a seed match and ≤5 AHC reads; green: subset of genes with a seed sequence for the indicated seed family and >5 reads in the AHC.

**Supplemental Figure 4 rel. to Figure 6**
**GSEA pathway analysis at 24h and 48h**
Naïve $CD4^+$ T cells from $T^{1792\Delta/\Delta}$, wt and $T^{1792tg/tg}$ mice were activated with plate-bound αCD28 and αCD3 mAbs for 0 and 24h. Total RNA was extracted for sequencing. Volcano plot of GSEA pathway (`KEGG,REACTOME,PID,BIOCARTA`) enrichment for genes differentially expressed after 24h (A) and 48h (B) activation between $T^{1792\Delta/\Delta}$ and $T^{1792tg/tg}$. Red dots indicate pathway enriched for genes up regulated whereas blue dots are pathways enriched for down regulated genes. Only significantly enriched pathways (1% FDR) are colored. Dot width indicates the gene set size for each pathway.

**Supplemental Figure 5 rel. to Figure 6**

**Transgenic miR-17~92 functionally corrected the defects of $CD28^{-/-}$ T cells to differentiate into $T_H1$, $T_H17$ and iTreg in vitro**

Naïve $CD4^+$ T cells were activated for 24h, 48h, and 72h with plate-bound αCD28 and αCD3 under in vitro skewing conditions for $T_H1$, $T_H17$, and $iT_{reg}$ differentiation. wt (black), $CD28^{-/-}$ (purple), rescue (dark blue) and $T^{1792tg/tg}$ (light blue). A) $T_H1$ differentiation assessed by IFNγ/Tbx21 staining in viable $CD4^+$ T cells B) $T_H17$ differentiation assessed by IL-17A/Rorγt staining in viable $CD4^+$ T cells C) $iT_{reg}$s characterized by CD25/ Foxp3 expression. Data from 2 independent experiments.

**Supplemental Figure 6 rel. to Figure 6**

**miR-17~92 expression modifies metabolism in activated $CD4^+$ T cells after 48h**

$CD4^+$ T cells from $T^{1792\Delta/\Delta}$ (grey), wt (black), $T^{1792tg/tg}$ (light blue) were assessed for their metabolic activity by mitochondria stress test with the seahorse machine. A) Mitochondria stress test measured with a 96-well seahorse in naïve $CD4^+$ T cells. 2 experiments with 3-4 biological replicates per group and experiment are shown. B) Basal respiration and ATP coupled respiration in naïve $CD4^+$ T cells C) Mitochondria stress test measured in $CD4^+$ T cells activated for 48h. D) Basal respiration and ATP coupled respiration in activated $CD4^+$ T cells. Pooled 3-8 biological replicates from 4 experiments are shown. Tukey's multiple comparison test, p values: *<0.05. E) RNA sequencing data shows an enrichment of genes associated with TCA. Shown is gene set REACTOME_TCA_CYCLE_AND_RESPIRATORY_ ELECTRON_TRANSPORT enrichment with genes differentially expressed between $T^{1792\Delta/\Delta}$ and $T^{1792tg/tg}$ in naïve, 24h and 48h post activation. Colors indicate fold change direction with blue being upregulated in $T^{1792\Delta/\Delta}$ and red downregulated.

**Supplemental Figure 7 rel. to Figure 7**

**All miR-17~92 seed families are necessary for CD28-induced gene repression**

Total RNA was extracted for sequencing from $CD4^+$ T cells. A) Correlation of gene expression levels between RNA-seq experiment 1 (Fig. 5) and 2 (Fig. 7). Black: genes without seed match, <5 AHC reads and no differential expression in the second RNA sequencing. Green: genes with a conserved binding site for each indicated seed family and >5 reads in the AHC; red: genes with a conserved binding site for each indicated seed family, >5 reads in the AHC and differential expression in the second RNA sequencing data set. B) For each seed family, dot plots compare the log2 value of the exonic expression ratio for each gene in the $CD28^{-/-}$ vs. wt comparison (x-axis) versus the exonic expression ratio for each gene in rescue vs. wt (y-axis) in $CD4^+$ T cells 24h post activation. Each ratio is compared to the cumulative fraction of all log2 ratios in the corresponding comparison. Black curve: all genes of our data set without a seed match and five or less AHC reads; green: subset of genes with a seed sequence for the indicated seed family (TS) and >5 reads in the AHC; red: genes with a conserved binding site for the corresponding seed family (TS), >5 reads in the AHC and differential expression in the second RNA sequencing data set.

**Table S1, related to Figure 5:** Doelz et al., The non-coding RNA miR-17~92 is a central mediator of T cell activation.

List of genes that fulfilled the following criteria: i) significant derepression in T1792Δ/Δ vs wt and significant repression in T1792tg/tg vs wt at 24h ii) predicted TS match iii) >5 AHC reads and iv) posttranscriptional regulation based on EISA. Applying these criteria across the 4 seed families from miR-17~92 cluster defined a set of 68 empirically supported direct miR-17~92 target genes. Individual seed families are reported.

| ENTREZID | SYMBOL | GENENAME | Length | miR |
|---|---|---|---|---|
| 11308 | Abi1 | abl-interactor 1 | 3505 | mmu-miR-106b-5p |
| 236511 | Ago1 | argonaute RISC catalytic subunit 1 | 7146 | mmu-miR-106b-5p |
| 70797 | Ankib1 | ankyrin repeat and IBR domain containing 1 | 7081 | mmu-miR-106b-5p |
| 433667 | Ankrd13c | ankyrin repeat domain 13c | 2654 | mmu-miR-106b-5p |
| 228359 | Arhgap1 | Rho GTPase activating protein 1 | 3326 | mmu-miR-106b-5p |
| 71704 | Arhgef3 | Rho guanine nucleotide exchange factor (GEF) 3 | 4585 | mmu-miR-106b-5p |
| 109168 | Atl3 | atlastin GTPase 3 | 6699 | mmu-miR-106b-5p |
| 20238 | Atxn1 | ataxin 1 | 10607 | mmu-miR-106b-5p |
| 12753 | Clock | circadian locomotor output cycles kaput | 10574 | mmu-miR-106b-5p |
| 231464 | Cnot6l | CCR4-NOT transcription complex, subunit 6-like | 9454 | mmu-miR-106b-5p |
| 74114 | Crot | carnitine O-octanoyltransferase | 4885 | mmu-miR-106b-5p |
| 98193 | Dcaf8 | DDB1 and CUL4 associated factor 8 | 7368 | mmu-miR-106b-5p |
| 75221 | Dpp3 | dipeptidylpeptidase 3 | 4940 | mmu-miR-106b-5p |
| 242960 | Fbxl5 | F-box and leucine-rich repeat protein 5 | 7479 | mmu-miR-106b-5p |
| 215751 | Ginm1 | glycoprotein integral membrane 1 | 1591 | mmu-miR-106b-5p |
| 83924 | Gpr137b | G protein-coupled receptor 137B | 3273 | mmu-miR-106b-5p |
| 73389 | Hbp1 | high mobility group box transcription factor 1 | 4753 | mmu-miR-106b-5p |
| 16451 | Jak1 | Janus kinase 1 | 5299 | mmu-miR-106b-5p |
| 231986 | Jazf1 | JAZF zinc finger 1 | 3098 | mmu-miR-106b-5p |
| 71819 | Kif23 | kinesin family member 23 | 5037 | mmu-miR-106b-5p |
| 17113 | M6pr | mannose-6-phosphate receptor, cation dependent | 2222 | mmu-miR-106b-5p |

| | | | | |
|---|---|---|---|---|
| 216001 | Micu1 | mitochondrial calcium uptake 1 | 7602 | mmu-miR-106b-5p |
| 54484 | Mkrn1 | makorin, ring finger protein, 1 | 3982 | mmu-miR-106b-5p |
| 56174 | Nagk | N-acetylglucosamine kinase | 1422 | mmu-miR-106b-5p |
| 69721 | Nkiras1 | NFKB inhibitor interacting Ras-like protein 1 | 4847 | mmu-miR-106b-5p |
| 192292 | Nrbp1 | nuclear receptor binding protein 1 | 2647 | mmu-miR-106b-5p |
| 244650 | Phlpp2 | PH domain and leucine rich repeat protein phosphatase 2 | 8388 | mmu-miR-106b-5p |
| 19344 | Rab5b | RAB5B, member RAS oncogene family | 4354 | mmu-miR-106b-5p |
| 19357 | Rad21 | RAD21 cohesin complex component | 3605 | mmu-miR-106b-5p |
| 53902 | Rcan3 | regulator of calcineurin 3 | 5062 | mmu-miR-106b-5p |
| 73469 | Rnf38 | ring finger protein 38 | 6831 | mmu-miR-106b-5p |
| 56613 | Rps6ka4 | ribosomal protein S6 kinase, polypeptide 4 | 3132 | mmu-miR-106b-5p |
| 54650 | Sfmbt1 | Scm-like with four mbt domains 1 | 8122 | mmu-miR-106b-5p |
| 20499 | Slc12a7 | solute carrier family 12, member 7 | 7313 | mmu-miR-106b-5p |
| 98396 | Slc41a1 | solute carrier family 41, member 1 | 4661 | mmu-miR-106b-5p |
| 58244 | Stx6 | syntaxin 6 | 3613 | mmu-miR-106b-5p |
| 69470 | Tmem127 | transmembrane protein 127 | 4401 | mmu-miR-106b-5p |
| 77975 | Tmem50b | transmembrane protein 50B | 2222 | mmu-miR-106b-5p |
| 100201 | Tmem64 | transmembrane protein 64 | 4681 | mmu-miR-106b-5p |
| 30934 | Tor1b | torsin family 1, member B | 3007 | mmu-miR-106b-5p |
| 216558 | Ugp2 | UDP-glucose pyrophosphorylase 2 | 4486 | mmu-miR-106b-5p |
| 75580 | Zbtb4 | zinc finger and BTB domain containing 4 | 7884 | mmu-miR-106b-5p |
| 232430 | Crebl2 | cAMP responsive element binding protein-like 2 | 2536 | mmu-miR-18a-5p |
| 75580 | Zbtb4 | zinc finger and BTB domain containing 4 | 7884 | mmu-miR-18a-5p |
| 11566 | Adss | adenylosuccinate synthetase, non muscle | 3570 | mmu-miR-19b-3p |

| | | | | |
|---|---|---|---|---|
| 236511 | Ago1 | argonaute RISC catalytic subunit 1 | 7146 | mmu-miR-19b-3p |
| 70797 | Ankib1 | ankyrin repeat and IBR domain containing 1 | 7081 | mmu-miR-19b-3p |
| 228359 | Arhgap1 | Rho GTPase activating protein 1 | 3326 | mmu-miR-19b-3p |
| 20238 | Atxn1 | ataxin 1 | 10607 | mmu-miR-19b-3p |
| 70533 | Btf3l4 | basic transcription factor 3-like 4 | 4265 | mmu-miR-19b-3p |
| 232196 | C87436 | expressed sequence C87436 | 3752 | mmu-miR-19b-3p |
| 12380 | Cast | calpastatin | 4747 | mmu-miR-19b-3p |
| 216527 | Ccm2 | cerebral cavernous malformation 2 | 3178 | mmu-miR-19b-3p |
| 12753 | Clock | circadian locomotor output cycles kaput | 10574 | mmu-miR-19b-3p |
| 104625 | Cnot6 | CCR4-NOT transcription complex, subunit 6 | 7561 | mmu-miR-19b-3p |
| 231464 | Cnot6l | CCR4-NOT transcription complex, subunit 6-like | 9454 | mmu-miR-19b-3p |
| 74256 | Cyld | CYLD lysine 63 deubiquitinase | 10569 | mmu-miR-19b-3p |
| 70186 | Fam162a | family with sequence similarity 162, member A | 635 | mmu-miR-19b-3p |
| 73389 | Hbp1 | high mobility group box transcription factor 1 | 4753 | mmu-miR-19b-3p |
| 231986 | Jazf1 | JAZF zinc finger 1 | 3098 | mmu-miR-19b-3p |
| 192292 | Nrbp1 | nuclear receptor binding protein 1 | 2647 | mmu-miR-19b-3p |
| 67229 | Prpf18 | pre-mRNA processing factor 18 | 3413 | mmu-miR-19b-3p |
| 19344 | Rab5b | RAB5B, member RAS oncogene family | 4354 | mmu-miR-19b-3p |
| 70510 | Rnf167 | ring finger protein 167 | 2078 | mmu-miR-19b-3p |
| 73469 | Rnf38 | ring finger protein 38 | 6831 | mmu-miR-19b-3p |
| 266781 | Snx17 | sorting nexin 17 | 1941 | mmu-miR-19b-3p |
| 229521 | Syt11 | synaptotagmin XI | 4993 | mmu-miR-19b-3p |
| 21815 | Tgif1 | TGFB-induced factor homeobox 1 | 2674 | mmu-miR-19b-3p |
| 100201 | Tmem64 | transmembrane protein 64 | 4681 | mmu-miR-19b-3p |
| 74868 | Tmem65 | transmembrane protein 65 | 3644 | mmu-miR-19b-3p |
| 30934 | Tor1b | torsin family 1, member B | 3007 | mmu-miR-19b-3p |
| 70827 | Trak2 | trafficking protein, kinesin binding 2 | 6240 | mmu-miR-19b-3p |
| 226757 | Wdr26 | WD repeat domain 26 | 7047 | mmu-miR-19b-3p |

| | | | | |
|---|---|---|---|---|
| 75580 | Zbtb4 | zinc finger and BTB domain containing 4 | 7884 | mmu-miR-19b-3p |
| 231570 | A830010M20Rik | RIKEN cDNA A830010M20 gene | 6756 | mmu-miR-92a-3p |
| 70797 | Ankib1 | ankyrin repeat and IBR domain containing 1 | 7081 | mmu-miR-92a-3p |
| 20238 | Atxn1 | ataxin 1 | 10607 | mmu-miR-92a-3p |
| 100383 | Bsdc1 | BSD domain containing 1 | 2659 | mmu-miR-92a-3p |
| 217946 | Cdca7l | cell division cycle associated 7 like | 2611 | mmu-miR-92a-3p |
| 98193 | Dcaf8 | DDB1 and CUL4 associated factor 8 | 7368 | mmu-miR-92a-3p |
| 57431 | Dnajc4 | DnaJ heat shock protein family (Hsp40) member C4 | 843 | mmu-miR-92a-3p |
| 76740 | Efr3a | EFR3 homolog A | 10045 | mmu-miR-92a-3p |
| 80517 | Herpud2 | HERPUD family member 2 | 2744 | mmu-miR-92a-3p |
| 338366 | Mia3 | melanoma inhibitory activity 3 | 7558 | mmu-miR-92a-3p |
| 17764 | Mtf1 | metal response element binding transcription factor 1 | 7956 | mmu-miR-92a-3p |
| 244650 | Phlpp2 | PH domain and leucine rich repeat protein phosphatase 2 | 8388 | mmu-miR-92a-3p |
| 19357 | Rad21 | RAD21 cohesin complex component | 3605 | mmu-miR-92a-3p |
| 73469 | Rnf38 | ring finger protein 38 | 6831 | mmu-miR-92a-3p |
| 56613 | Rps6ka4 | ribosomal protein S6 kinase, polypeptide 4 | 3132 | mmu-miR-92a-3p |
| 75627 | Snapc1 | small nuclear RNA activating complex, polypeptide 1 | 2125 | mmu-miR-92a-3p |
| 21815 | Tgif1 | TGFB-induced factor homeobox 1 | 2674 | mmu-miR-92a-3p |
| 70827 | Trak2 | trafficking protein, kinesin binding 2 | 6240 | mmu-miR-92a-3p |
| 216558 | Ugp2 | UDP-glucose pyrophosphorylase 2 | 4486 | mmu-miR-92a-3p |

**Table S2, related to Figure 7:** Doelz et al., The non-coding RNA miR-17~92 is a central mediator of T cell activation.

List of genes in which “rescue” T cells were more similar to wt and $T^{1792tg/tg}$ cells than $CD28^{-/-}$ and $T^{1792\Delta/\Delta}$ cells. Corresponds to genes in box in figure 7D.

| Gene name |
|---|
| Tgtp2 |
| Gbp6 |
| Mir6974 |
| Oaf |
| Smim3 |
| AW112010 |
| Coro2a |
| Slco3a1 |
| Tspan9 |
| Ly6a |
| H2-Q6 |
| Slc13a3 |
| Atf3 |
| Cacnb3 |
| Casp1 |
| Casp4 |
| Runx2 |
| Cd44 |
| Socs1 |
| Cxcr3 |
| Ccr5 |
| Eomes |
| H2-T23 |
| Hbegf |
| Icam1 |
| Irgm1 |
| Cxcl10 |
| Ifng |
| Igtp |
| Il12a |
| Il12rb2 |
| Il4 |
| Ajuba |
| Cxcl9 |
| Gbp4 |
| Penk |
| Ppic |
| Ptger3 |
| Ell2 |
| Rab20 |

Rpl29
Sox5
Serpinb6b
Serpina3g
Serpinb9
Stra6
Tcf4
Fam26f
Tgtp1
Soat2
Xlr3c
Gm4841
C1ql3
Ppp1r16b
Gbp7
Asap3
Picalm
Ccdc102a
Serpina3f
Gadd45g
Mgll
Gm4951
Ubd
Ehd2
Insl6
Batf3
Gm996
Rnf225
Brsk1
Mir18
Mir20a
AA467197
Dlc1
Irgm2
Gbp3
Espn
Ube2l6
Ly6i
Iigp1
Il21
Gm12185
Gbp10
Gm12250
4930428O21Rik
Fkbp11
Tma7
Arrdc4
Tmem35a

Fam212a
Tln2
Zbtb46
Mir17
Nuak2
Rnf19b
Abhd14b
Sytl3
Trps1
Tnfrsf25
Gm1966
Gm8989
Il10ra
Zbp1
Gbp8
Tgfbr3
Nckap5l
Gprin3
Gm18853
Gvin1
Ptprn
Parp3
Cysltr2
H2-DMa

**Supplementary Table I**
List of genes that fulfilled the following criteria: i) significant derepression in $T^{1792\Delta/\Delta}$ vs wt and significant repression in $T^{1792tg/tg}$ vs wt at 24h ii) predicted TS match iii) >5 AHC reads and iv) posttranscriptional regulation based on EISA. Applying these criteria across the 4 seed families from miR-17~92 cluster defined a set of 68 empirically supported direct miR-17~92 target genes. Individual seed families are reported.

**Supplementary Table II**
List of genes in which "rescue" T cells were more similar to wt and $T^{1792tg/tg}$ cells than $CD28^{-/-}$ and $T^{1792\Delta/\Delta}$ cells. Corresponds to genes in box in figure 7D.
